# Intratumoral mregDC and CXCL13 T helper niches enable local differentiation of CD8 T cells following PD-1 blockade

**DOI:** 10.1101/2022.06.22.497216

**Authors:** Assaf Magen, Pauline Hamon, Nathalie Fiaschi, Leanna Troncoso, Etienne Humblin, Darwin D’souza, Travis Dawson, Matthew D. Park, Joel Kim, Steven Hamel, Mark Buckup, Christie Chang, Alexandra Tabachnikova, Hara Schwartz, Nausicaa Malissen, Yonit Lavin, Alessandra Soares-Schanoski, Bruno Giotti, Samarth Hegde, Raphaël Mattiuz, Clotilde Hennequin, Jessica Le Berichel, Zhen Zhao, Stephen Ward, Isabel Fiel, Colles Price, Nicolas Fernandez, Jiang He, Baijun Kou, Michael Dobosz, Lianjie Li, Christina Adler, Min Ni, Yi Wei, Wei Wang, Namita T. Gupta, Kunal Kundu, Kamil Cygan, Raquel P. Deering, Alex Tsankov, Seunghee Kim-Schulze, Sacha Gnjatic, Ephraim Kenigsberg, Myron Schwartz, Thomas U. Marron, Gavin Thurston, Alice O. Kamphorst, Miriam Merad

**Author notes:** These authors contributed equally (Assaf Magen, Pauline Hamon).

## Abstract

Here, we leveraged a large neoadjuvant PD-1 blockade trial in patients with hepatocellular carcinoma (HCC) to search for correlates of response to immune checkpoint blockade (ICB) within T cell-rich tumors. We show that ICB response correlated with the clonal expansion of intratumoral CXCL13^+^ CH25H^+^ IL-21^+^ PD-1^+^ CD4 T helper cells (CXCL13^+^ Th) and Granzyme K^+^ PD-1^+^ effector-like CD8 T cells, whereas terminally exhausted CD39^hi^ TOX^hi^ PD-1^hi^ CD8 T cells dominated in non-responders. Strikingly, most T cell receptor (TCR) clones that expanded post-treatment were found in pre-treatment biopsies. Notably, PD-1^+^ TCF-1^+^ progenitor-like CD8 T cells were present in tumors of responders and non-responders and shared clones mainly with effector-like cells in responders or terminally differentiated cells in non-responders, suggesting that local CD8 T cell differentiation occurs upon ICB. We found that these progenitor CD8 T cells interact with CXCL13^+^ Th cells within cellular triads around dendritic cells enriched in maturation and regulatory molecules, or “mregDC”. Receptor-ligand analysis revealed unique interactions within these triads that may promote the differentiation of progenitor CD8 T cells into effector-like cells upon ICB. These results suggest that discrete intratumoral niches that include mregDC and CXCL13^+^ Th cells control the differentiation of tumor-specific progenitor CD8 T cell clones in patients treated with ICB.

## Introduction

Surgical resection is the preferred treatment of early hepatocellular carcinoma (HCC) lesions, but more than half of these tumors recur within two years (Roayaie et al., 2013; Tabrizian et al., 2015), presumably due to residual micrometastatic diseases (Shindoh et al., 2013). These trends highlight the need for perioperative therapy to improve HCC outcomes. Neoadjuvant immune checkpoint blockade (ICB) targeting the PD-1/PD-L1 axis has been successful in inducing pathological response and preventing recurrence in multiple tumor types, in part by driving the expansion of tumor-specific T cells, which may also induce systemic immunity and eliminate micrometastases (Chalabi et al., 2020; Forde et al., 2018; Huang et al., 2019; Marron et al., 2022a, 2022b; Topalian et al., 2020). Neoadjuvant trials also leverage the unique advantage of permitting extensive molecular characterization of treated surgical resections, thereby enabling us to query mechanisms of response or resistance to immunotherapy (Marron et al., 2022b). We recently led a neoadjuvant clinical trial for early-stage HCC patients, in which treatment-naive patients received two doses of PD-1 blockade prior to surgery (Marron et al., 2022a). We observed a 30% pathological response rate, which prompted a more detailed investigation into the cellular and molecular pathways that promote an effective anti-tumor immune response.

T cell infiltration is a well-established prognostic factor of ICB response (Chen and Mellman, 2017; Li et al., 2021), and generally, three main patterns of T cell infiltration have been described in tumor lesions (Galon et al., 2006; Gruosso et al., 2019): (1) tumors with high T cell content in the tumor core (referred to hereafter as “T cell rich”), (2) tumors in which T cell infiltration is restricted to the stroma (“T cell excluded”), and (3) tumors with overall low T cell content (“T cell low”). The T cell rich infiltration pattern is the most conducive to ICB response, though it is not an accurate predictor of response (Galon and Bruni, 2019).

Response to ICB has been associated with an increase in tumor infiltrating PD-1^hi^ CD8 T cells in several clinical studies (Bassez et al., 2021; Huang et al., 2019; Tumeh et al., 2014). Most recently, PD-1^hi^ CD8 T cells expressing intermediate levels of checkpoint molecules (PD-1, LAG-3, CTLA-4) and high levels of effector molecules (Granzyme K) were associated with a potent response to a combination of PD-1 blockade and chemotherapy in non-small cell lung cancer (Liu et al., 2022). However, it remains unclear whether the induction of an effective anti-tumor CD8 T cell response occurs primarily at the local tumor microenvironment (TME) or in tumor-draining lymph nodes (tdLN) (van der Leun et al., 2020; Liu et al., 2022; Wu et al., 2020; Yost et al., 2019). It is also unclear whether antigen-experienced CD8 T cells need to be reactivated by antigen-presenting cells (APC), including macrophages (Macs) and dendritic cells (DC), to respond to ICB. In addition to CD8 T cells, B cells (Cabrita et al., 2020; Helmink et al., 2020) and CXC chemokine ligand 13 (CXCL13)-expressing CD4 T cells have been associated with response to ICB (Liu et al., 2022), but how these cell types contribute to anti-tumor immunity remains elusive.

To probe the mechanisms of response to ICB in early-stage HCC, we analyzed surgically-resected tumor lesions and matched, non-involved liver tissues from patients who were responsive or resistant to ICB therapy. Not surprisingly, all tumor lesions from responsive patients were highly infiltrated by T cells; however, many non-responsive lesions were also enriched in T cells. Using multiplex imaging as well as paired single-cell RNA sequencing (scRNAseq) and single-cell T cell receptor sequencing (scTCRseq) of nearly one million immune cells isolated from tumor and adjacent non-involved tissues, we found that pathological response to ICB strongly correlates with the intratumoral expansion of PD-1^hi^ effector-like CD8 T cells and CD4 T cells expressing features of follicular helper T cells such as CXCL13 and IL-21 (referred to hereafter as CXCL13^+^ Th). scTCRseq analysis showed that these PD-1^hi^ effector-like CD8 T cells, but also terminally dysfunctional CD8 T cells, are two potential outcomes of a particular clonotype that enters a proliferative and progenitor-exhausted CD8 T cell state (Tpex, referred to hereafter as progenitor CD8 T cells), illustrating a bifurcation in the differentiation potential. Strikingly, an evaluation of the clonal distribution of TCRs across adjacent liver tissue, tdLN, peripheral blood mononuclear cells (PBMC), and pre-treatment tumor biopsies revealed that many T cell clones that had expanded in the tumor post-PD-1 blockade were already present at the tumor site prior to treatment. Finally, we show that interactions between progenitor CD8 T cells and CXCL13^+^ Th cells

occur within cellular triads around mregDC, mature DC that have entered a unique molecular state triggered upon capture and sensing of tumor antigens (Maier et al., 2020). Altogether, our results suggest that discrete intratumoral cellular niches, comprised of mregDC and CXCL13^+^ Th cells, enable the reactivation of pre-existing T cell clones into effective anti-tumor CD8 T cells.

## RESULTS

### A subset of T cell-rich tumors failed to respond to PD-1 blockade

HCC is prototypically an inflammation-driven cancer that develops after years of inflammatory insults. To examine the T cell distribution in HCC lesions from patients who either responded or resisted to ICB, we analyzed 29 early-stage HCC lesions and patient-matched non-involved liver specimens that were surgically resected after two doses of cemiplimab (20 patients, administered as part of our clinical trial NCT03916627) or 2-4 doses of nivolumab (9 patients, administered off-label). 30% (cemiplimab-treated) and 22% (nivolumab-treated) of patients across all HCC etiologies responded to ICB, which was defined as ≥50% tumor necrosis by pathological examination of resected lesions, based on an exploratory cutoff from our recent trial (Marron et al., 2022a) **(Figure S1A, Table S1)**. Immunohistochemistry (IHC) and immunofluorescence (IF) of resected tumor lesions showed high variability in the extent and distribution of T cell infiltrates, consistent with patterns previously described **(Figure 1A)** (Galon et al., 2006; Gruosso et al., 2019). All treatment-responsive lesions and 40% of non-responsive lesions were highly infiltrated with T cells **(Figure 1A-C)**. Immune aggregates – defined as regions with high densities of lymphocytes – were increased both in size and number in T cell rich lesions amongst non-responders, but were highest in responders **(Figure S1B)**. T cell infiltration and tumor response to ICB were not correlated with tumor mutational burden (TMB) **(Figure 1D)**, though *p53* (TP53) mutations were enriched in responders (P<0.001; Hypergeometric test), while β-catenin (CTNNB1) activated mutations were enriched in T cell low lesions **(Figure 1E)** (P=0.001), consistent with recent finding showing that WNT pathway activating mutations suppress immune cell infiltration in tumors (Ruiz de Galarreta et al., 2019; Spranger et al., 2015).

**Figure 1.**
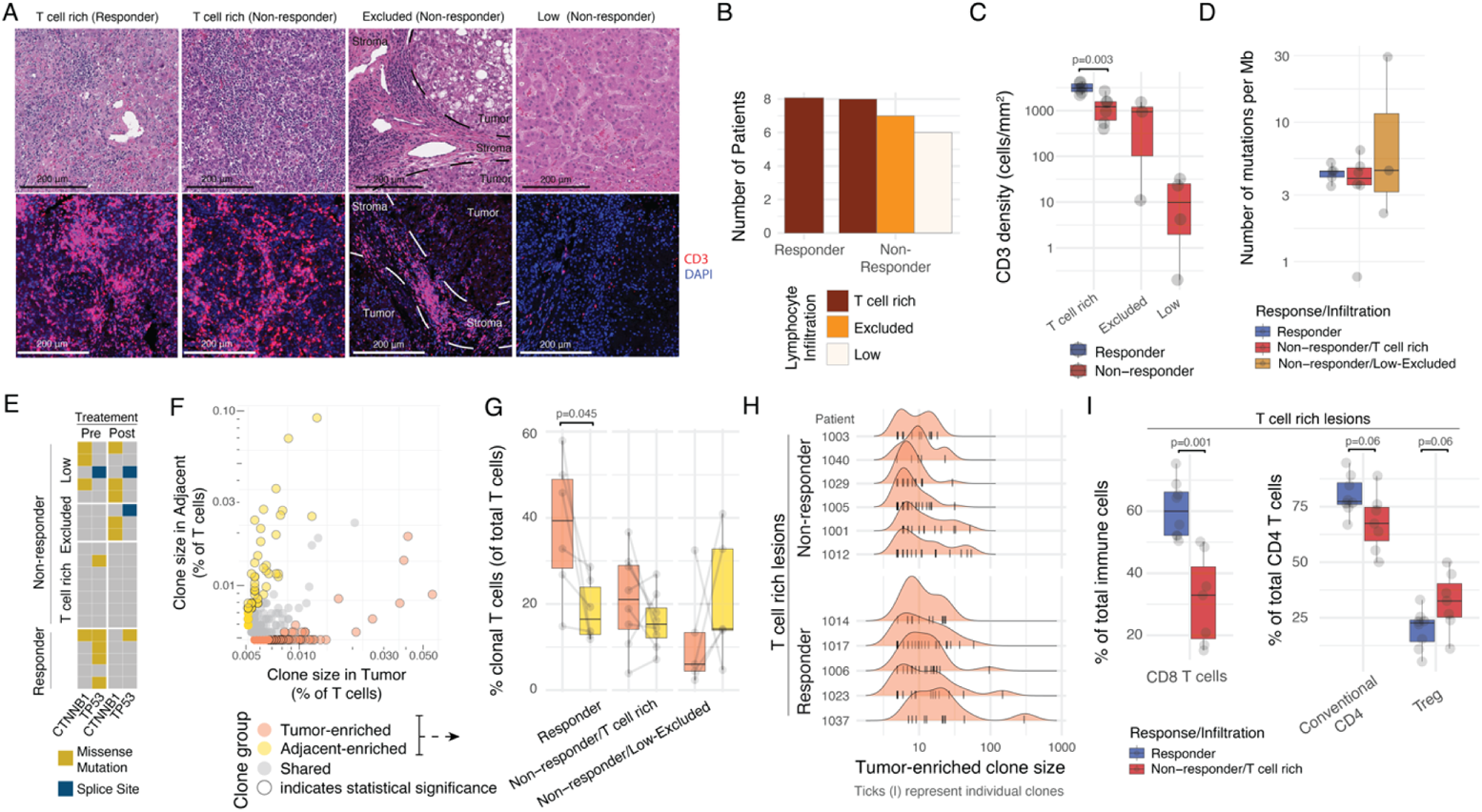
A subset of T cell rich tumors failed to respond to PD-1 blockade. Surgically resected HCC lesions were isolated after two or more doses of PD-1 blockade and analyzed by H&E (N=20), immunofluorescence (IF, N=20), single-cell RNA and TCR sequencing (scRNAseq and scTCRseq, N=29 and N=21, respectively), and whole-exome sequencing (WES, N=20). HCC tumor biopsies (N=20) were collected prior to neoadjuvant PD-1 blockade and analyzed by WES. (**A-C**) Assessment of spatial distribution patterns in HCC by H&E and IF. (A) H&E (top) and CD3 IF (bottom) of representative tumor lesions across distinct immune infiltration patterns. (B) Distribution of immune infiltration pattern across responders and non-responders. (C) Distribution of CD3^+^ T cell density by IF across responders and non-responders stratified by immune infiltration pattern. (**D-E**) Mutational analysis of tumor lesions using WES. (D) Tumor mutation burden (TMB) quantification across responders and non-responders stratified by immune infiltration pattern. (E) Mutational status of β-catenin (CTNNB1) and p53 (TP53) across patient groups, pre and post therapy. (**F-I**) scRNAseq and scTCRseq analysis of T cell clonality across the tumor and adjacent tissues. (F) Frequencies and classification of unique TCRs observed by scTCRseq in tumor (x axis) or adjacent (y axis) in a representative sample. (G) Frequencies of clonal T cells mapping to tumor- and adjacent-enriched from (F), stratified by response and immune infiltration pattern. (H) Histograms of clone size (number of cells per clone) distribution in tumor per patient, stratified by response and immune infiltration pattern. Ticks represent individual clones. (I) Cellular abundances by scRNAseq for CD8 T cells (left) and conventional and regulatory CD4 T cells (right) across responders and T cell rich non-responders.

We then used paired scRNAseq/scTCRseq to characterize the molecular profile of 918,811 CD45^+^ cells across tumors and adjacent tissues. We also searched for and characterized T cells that shared the same TCR (clonal T cells) **(Figure 1F)**, while T cells with no other identical TCR were termed singlets. In responders, clonal T cells were significantly more abundant in tumor tissues compared to adjacent tissues and 40% of T cells mapped to tumor-enriched clones, the largest of which consisted of 300 cells **(Figure 1G-H)**. In contrast, clonal T cells were only slightly more enriched in tumors compared to adjacent tissues, in T cell rich non-responders **(Figure 1G)**.

Clustering analysis of the scRNAseq data identified 107 clusters of immune cells, which we segregated into CD8 (33 clusters), conventional CD4 (17 clusters) and regulatory CD4 (Tregs) (5 clusters), naive and proliferating T cell clusters which included both CD8 from CD4 T cells, as well as other lymphoid and myeloid populations **(Figure S1C, Table S2)**. Consistent with our reports in NSCLC (Lavin et al., 2017; Leader et al., 2021), conventional CD4 T cells and Tregs were highly enriched in tumors compared to adjacent tissues, whereas NK cells and CD16^+^ monocytes were reduced in the TME **(Figure S1D)**. CD8 T cells – the most abundant immune cell type – were significantly increased in responders, when compared to T cell rich non-responders **(Figure 1I)**. Conventional CD4 T cells were more abundant in responders, whereas Tregs were more abundant in T cell rich non-responders.

### Identification of distinct molecular phenotypes among tumor-infiltrating CD8 and CD4 T cells in HCC patients that respond to PD-1 blockade

Given that the molecular mechanisms that promote tumor resistance to ICB are likely very distinct in T cell rich versus T cell low or excluded lesions, we sought to focus our analysis primarily on T cell rich tumor lesions. In addition to cellular clusters, we defined groups of co-expressed genes associated with effector, memory and cytotoxic features **(Figure S2A-B)**. CD8 T cell clusters were pooled into 7 clusters based on such similarities **(Figure S2A)**; four of these (1-4) highly expressed molecules features of chronic antigen activation and exhaustion such as *PDCD1* (PD-1), *CTLA4*, *TOX2*, *HAVCR2* (TIM-3) **(Figure 2A-B)**, as well as high levels of *CXCL13* and *DUSP4* (referred to hereafter as PD-1^hi^ CD8 T cells). The remaining three clusters (5-7) showed reduced features of chronic antigen stimulation but higher levels of AP-1 expression (*FOSB*). These PD-1^lo^ CD8 T cells were organized according to the co-expression of gene modules associated with effector function (*GZMK*, cluster 5), memory (*TCF7*, *LEF1*, cluster 6) and cytotoxicity (*GNLY* and *TYROBP*, cluster 7). All four PD-1^hi^CD8 T cell clusters were highly enriched in the TME, whereas PD-1^lo^ CD8 T cells were enriched in adjacent tissues. In accordance with previous reports (Gros et al., 2014; Hanada et al., 2022; Simoni et al., 2018) and tumor-enrichment, we propose that PD-1^hi^ CD8 T cells constitute tumor-specific T cells **(Figure 2C).**

**Figure 2.**
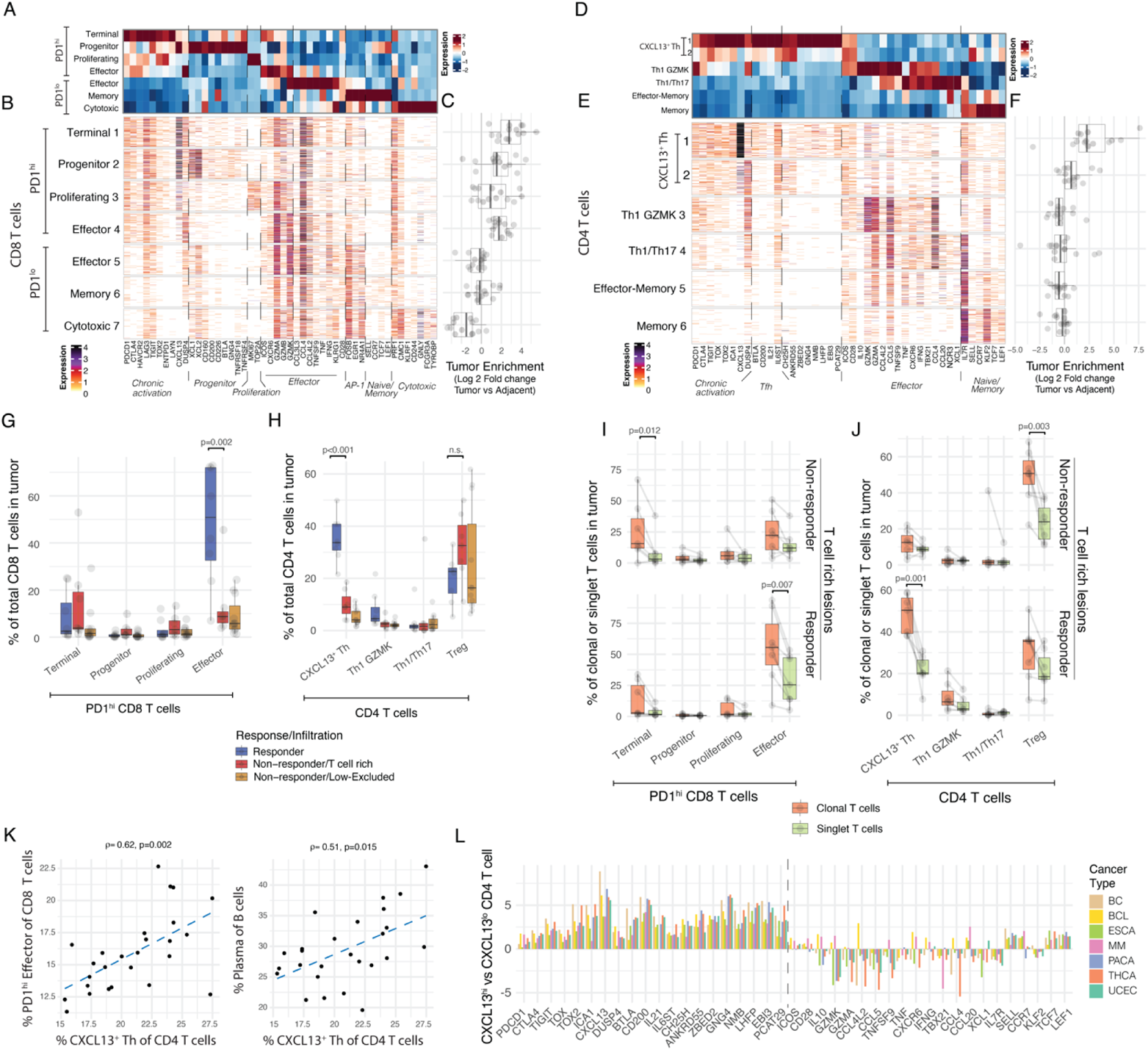
Responders are characterized by a distinct molecular phenotype of CD8 and CD4 T cells clonally expanded in a tumor specific manner. (**A-C**) Expression profiling of CD8 T cell cluster-defining genes by scRNAseq (A) showing column standardized average expression and (B) number of UMI per cell and (C) CD8 cluster frequencies enrichment in tumor versus adjacent tissue. (**D-F**) Expression profiling of CD4 T cell cluster-defining genes by scRNAseq (D) showing column standardized average expression and (E) number of UMI per cell and (F) CD4 cluster frequencies enrichment in tumor versus adjacent tissue. (**G-H**) Cluster frequencies stratified by response and immune infiltration pattern in tumor samples for (G) PD-1^hi^ CD8 T cells and (H) CD4 T cells. (**I-J**) Cluster frequencies among tumor-enriched and tumor singlet clones stratified by response and immune infiltration pattern in tumor samples for key (I) CD8 T cells and (J) CD4 T cell clusters. (**K**) Correlation between cellular abundances of CXCL13^+^ Th cells to PD-1^hi^ Effector CD8 T and B cells by scRNAseq. (**L**) Analysis of CXCL13^+^ Th transcriptional pattern across cancer types from external datasets. Showing log 2 Fold change between CXCL13^hi^ Th and non-CXCL13^lo^ Th cells, for CD4 molecules from (D-E) by scRNAseq (BC – breast cancer, BCL – B-cell lymphoma, MM – multiple myeloma, PACA – pancreatic cancer, UCEC – uterine corpus endometrial carcinoma, THCA – thyroid carcinoma, ESCA – esophageal cancer).

Among PD-1^hi^ CD8 T cells, the first cluster expressed higher levels of inhibitory molecules (*PDCD1*, *CTLA4*, *HAVCR2*, *TOX2*, *ENTPD1* (CD39)) and lower levels of effector and cytotoxic genes (*GZMK*, *CCL3L3*, *IFNG*, *KLRG1*), in line with a terminally-differentiated, exhausted state (thereby termed PD-1^hi^ Terminal). Cluster 2 expressed the highest levels of *XCL1*, as well as some expression of genes associated with naive/memory (*TCF7*, *LEF1*) and Tfh (*CD200*, *BTLA*, *GNG4*) features, the combination of which were characteristic of progenitor-exhausted CD8 T cells (PD-1^hi^ Progenitor, also called Tpex) (Eberhardt et al., 2021; Im et al., 2016). Cluster 3 (PD-1^hi^ Proliferating) resembled proliferating cells, given the expression of *MKI67*. Cluster 4 expressed lower levels of chronic activation features and higher levels of effector molecules including *GZMK* and *GNLY*, in line with an effector-like state (PD-1^hi^ Effector).

Similar analyses of the conventional CD4 T cell compartment revealed two clusters that displayed features of chronic activation and exhaustion (*PDCD1*, *LAG3*, *CTLA4*, *TOX2*) and features of follicular helper T cells (Tfh) including *BTLA*, *CD200*, *IL21* and *TCF7* (Crotty, 2014; Xin et al., 2015; Zander et al., 2019, 2022), as well as *IL6ST*, encoding the IL-6 receptor signal transducer, which is known to instruct Tfh cell differentiation and acts upstream of IL-21 (Crotty, 2014; Harker et al., 2013), and CXCL13. Notably, the latter molecules reflecting Tfh qualities have been shown to promote the recruitment of CXCR5-expressing B cells and possibly progenitor CD8 T cells (Brummelman et al., 2018; Im et al., 2016; Siddiqui et al., 2019) **(Figure 2D-E)**. As these clusters also expressed Th1 features such as *IFNG*, we designated these clusters as CXCL13^+^ helper T cells (CXCL13^+^ Th). Interestingly, CXCL13^+^ Th also expressed cholesterol 25-hydroxylase (*CH25H*) – the enzyme responsible for generating 7α,25-dihydroxycholesterol, the oxysterol chemoattractant for Epstein-Barr virus-induced G-protein coupled receptor 2 (EBI2, also known as GPR183) expressed on DC, monocyte-derived cells, and progenitor CD8 T cells (Eberhardt et al., 2021; Hudson et al., 2019). We also identified Th1-like cells that express *IFNG* and *GZMK*, as well as Th17-like cells, expressing *CCL20* and *NCR3*. The remaining CD4 T cell clusters expressed naive-memory programs (*IL7R*, *TCF7*, *CCR7* and *LEF1*), and some co-expressed effector genes (*GZMK*, *GZMA*, *IFNG*). Consistent with recent studies showing that CXCL13^+^ CD4 T cells are enriched for tumor-specific CD4 T cells (Hanada et al., 2022; Veatch et al., 2022), only CXCL13^+^ Th were significantly enriched in the TME compared to adjacent tissues **(Figure 2F)**. These data suggest that in concert with PD-1^hi^ CD8 T cells, CXCL13^+^ Th might play a more prominent role in anti-tumor immunity.

We also measured the prevalence of CD4 and CD8 T cell subsets in responders and non-responders. We found that both PD-1^hi^ effector-like CD8 T cells and CXCL13^+^ Th cells were significantly enriched in responders compared to non-responders **(Figure 2G-H, S2C-D)**. PD-1^hi^ effector CD8 T cells and CXCL13^+^ Th cells were also clonally expanded preferentially in tumors of responders **(Figure 2I-J)**, whereas PD-1^hi^ Terminal CD8 T cells and Tregs were clonally expanded preferentially in tumors of non-responders **(Figure 2I-J)**. In contrast, PD-1^lo^ CD8 T cells and effector/memory CD4 T cells were not detected amongst tumor-enriched clones altogether **(Figure S2E-F)**, suggesting that they might not be involved in the T cell response to tumor antigens. Notably, we found that CXCL13^+^ Th cell abundance was correlated with that of PD-1^hi^ effector CD8 T cells and plasma cells **(Figure 2K),** supporting the role of helper CD4 T cells in the differentiation of PD-1^+^ CD8 T cells towards an effector-like state (Zander et al., 2019, 2022) and in a multicellular response to ICB, alongside plasma cells (Patil et al., 2022).

PD-1^hi^ effector CD8 T cells indeed have been identified in multiple human cancers, but the classification and nomenclature of T cell subsets are quite variable across studies (van der Leun et al., 2020). Likewise, CXCL13^+^ CD4 T cells have also been recently described in several cancer studies (Liu et al., 2022; Noël et al., 2021; Zheng et al., 2021), yet the CXCL13^+^ Th program has not been extensively studied. To determine whether the CXCL13^+^ Th molecular program enriched in responders represents a conserved cell state, we probed seven published human cancer scRNAseq datasets (Zheng et al., 2021) and found *CXCL13*, *IL21*, *CH25H* as well as *TCF7* to be consistently over-expressed in CXCL13^hi^ CD4 T cells across cancer types **(Figure 2L)**. These results, together with the collective group of CD8 T cell molecular programs described in recent studies (van der Leun et al., 2020), suggest that CXCL13^+^ Th program could serve as a proxy of tumor reactive-T cells across multiple cancer types.

### CD8 and CD4 T cells expand and differentiate locally in the TME upon PD-1 blockade

To better understand the relationships among distinct CD8 T cell transcriptional states, we characterized the molecular composition of the top 10 tumor-enriched, expanded T cell clones in each patient. We found that most TCR clones encompassed the four major PD-1^hi^ CD8 subsets, indicating a local differentiation and expansion of CD8 T cells in tumor tissues of both responders and non-responders **(Figure 3A-B, S3A-B)**. Still, CD8 T cell differentiation was significantly skewed towards PD-1^hi^ effector-like cells in responders, whereas in non-responders, differentiation favored terminally-differentiated (PD-1^hi^ terminal) CD8 T cells **(Figure 3A-3B)**. We confirmed that the transcriptional patterns of cells sharing clones indeed corresponded to each of the four PD-1^hi^ CD8 molecular states in a representative responder **(Figure S3C)**. These results align with studies in mouse models that show PD-1^hi^ progenitors differentiate into effector-like and terminally-differentiated T cells (Hudson et al., 2019; Im et al., 2016; Miller et al., 2019; Siddiqui et al., 2019; Utzschneider et al., 2016).

**Figure 3.**
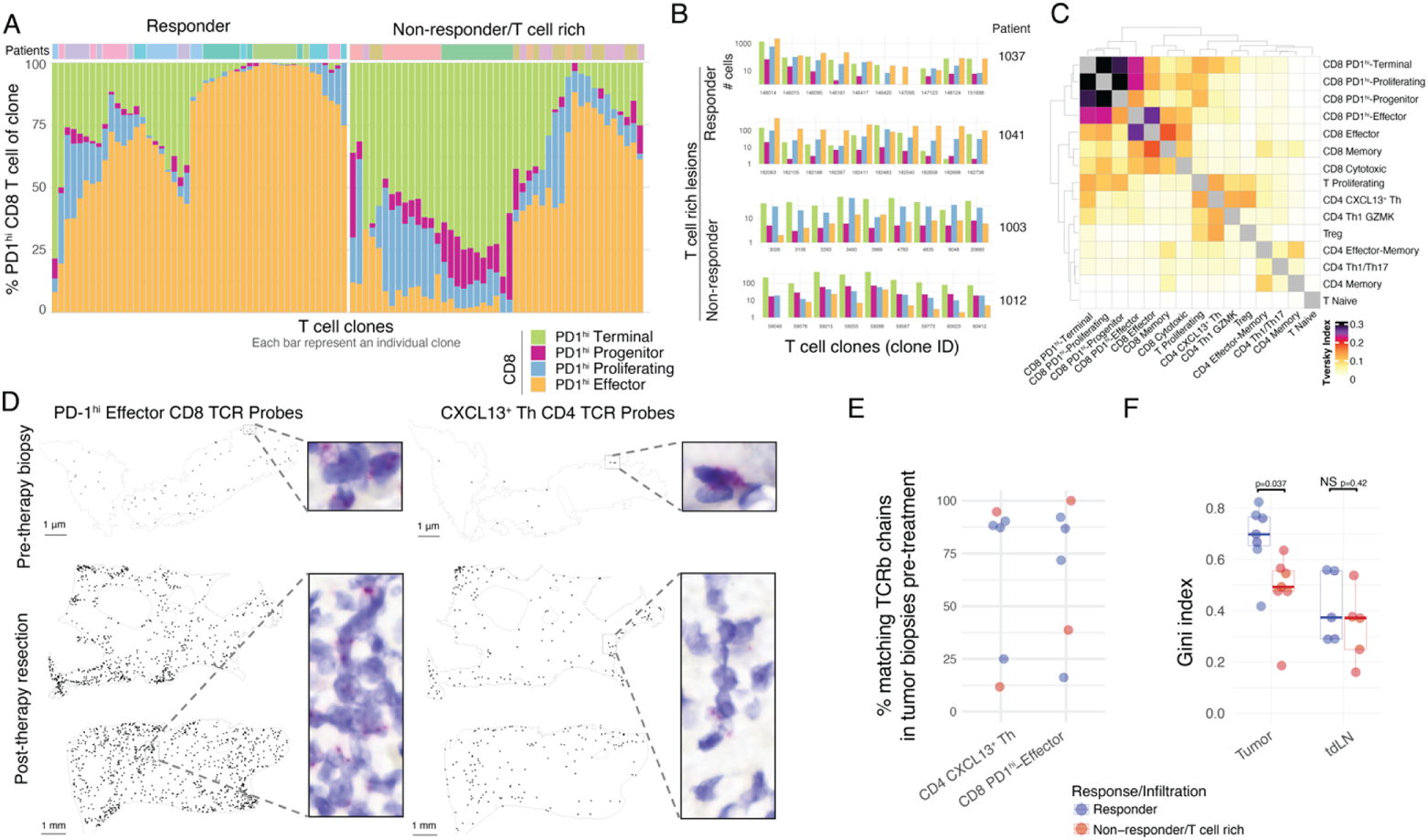
Local expansion of CD4 and CD8 T cells in the tumor upon PD-1 blockade. (**A-C**) Phenotypic analysis of clonotype sharing by scTCRseq. (A) Phenotypic distribution of PD-1^hi^ CD8 T cells in individual tumor-enriched clones (top 10 per patient), separately for responders and T cell rich non-responders. (B) Highlight of CD8 phenotypic distribution for selected CD8 clones from selected patients. (C) Trevsky index of TCR sharing across T cell clusters in tumor. (**D**) BaseScope TCR imaging analysis of 8 selected tumor-enriched clones in a responder patient (1006). Spatial distribution of 4 CD8 T cell and 4 CD4 T cell clones across biopsy and resection sample. (**E-F**) Pre-treatment biopsies analyzed by Bulk TCR sequencing, and scTCRseq of tdLN from time of resection. (**E**) Percent of post-treatment tumor-enriched clones present in pre-treatment tumor lesions by Bulk TCRseq and tdLN by scTCRseq, across responders and T cell rich non-responders. (**F**) Gini inequality index measure for T cell clonal expansion for clonal expansion in tumor and tdLN across responders and T cell rich non-responders.

By quantifying the Tversky asymmetric similarity index (Tversky, 1977), we further confirmed that PD-1^hi^ CD8 T cell clusters have a high degree of TCR sharing **(Figure 3C)**. We also identified substantial TCR sharing between CXCL13^+^ Th and a cluster of proliferating T cells **(Figure 3C, S1C)**, suggesting that like the PD-1^hi^ CD8 T cell clusters, CXCL13^+^ Th undergo a proliferative burst upon PD-1 blockade. To determine whether tumor-enriched T cell clones that expanded upon PD-1 blockade were present in the tumor prior to therapy – or alternatively, were expanded in the periphery and recruited upon treatment – we applied TCR imaging and TCRseq to analyze paired post-treatment tumor samples and pre-treatment biopsies. BaseScope analysis of TCRs corresponding to the top four tumor-enriched expanded clones of PD-1^hi^ effector-like CD8 T cells and CXCL13^+^ Th cells from a responder patient (patient 1006) revealed that these clones were present in the tumor prior to treatment, relative to the baseline signal reflecting negative control probes **(Figure 3D, S3D).** We also used bulk TCR sequencing of pre-treatment biopsies to search for TCRβ sequences that matched tumor-expanded CXCL13^+^ Th and PD-1^hi^ effector-like CD8 T cells from resected samples. In most patients, the majority of the TCRβ clones associated with CXCL13^+^ Th and PD-1^hi^ effector CD8 T cells were, in fact, present in the tumor prior to treatment **(Figure 3E, S3F)**.

To explore whether the expansion of tumor clones could also occur in tdLN upon PD-1 blockade, we examined T cell clones in tdLN samples to determine whether tumor-enriched clones were present (identified based on liver lymphatic drainage patterns) (Frenkel et al., 2020) collected on the day of resection, as well as in patient-matched PBMC. While a minority of shared clones were found in tdLN and blood with more matches in responders **(Figure S3F-G)**, T cell clonal expansion analysis (Gini inequality index, Methods) further revealed that tumor-expanded T cell clones were similarly expanded in the tdLN of responders and T cell rich non-responders **(Figure 3F, S3G)**. In contrast, tumor-expanded T cell clones were significantly more expanded in the TME of responders relative to T-cell-rich non-responders.

### Identification of cellular triads associated with the reactivation of PD-1^hi^ CD8 T cells in the TME

The observation that tumor clones expand and differentiate locally in responders led us to search for local cues that could contribute to the differentiation and/or maintenance of PD-1^hi^ effector-like CD8 T cells in responders. Because DC excel at instructing T cells, we hypothesized that specific DC programs may be responsible for coordinating the expansion of effector PD-1^hi^ CD8 T cells. Among the mononuclear phagocytes, the DC compartment includes cDC1 (*CLEC9A*), cDC2 (*CD1C*) and DC enriched in maturation and regulatory markers “mregDC” (*CD274*, *CCR7*, *CCL22*, *BIRC3*, *IDO1*, *IL4I1*, and notably *LAMP3*, which encodes DC-LAMP) **(Figure 4A)**, a molecular state induced in cDC upon capture of tumor antigens (Maier et al., 2020). We confirmed at the protein level that mregDC expressed the highest levels of co-stimulatory and inhibitory molecules (CD80/CD86, PD-L1 and PD-L2) amongst all other DC **(Figure S4A)**. mregDC also expressed the highest levels of MHC-II, consistent with our recent report that mregDC preferentially engage CD4 T cells in treatment-naive human NSCLC lesions (Cohen et al., 2022), and this motivated our exploration of potential mregDC-T cell interactions in HCC lesions that might underlie the local expansion of CXCL13^+^ Th and PD-1^hi^ effector-like CD8 T cells observed in responders.

**Figure 4.**
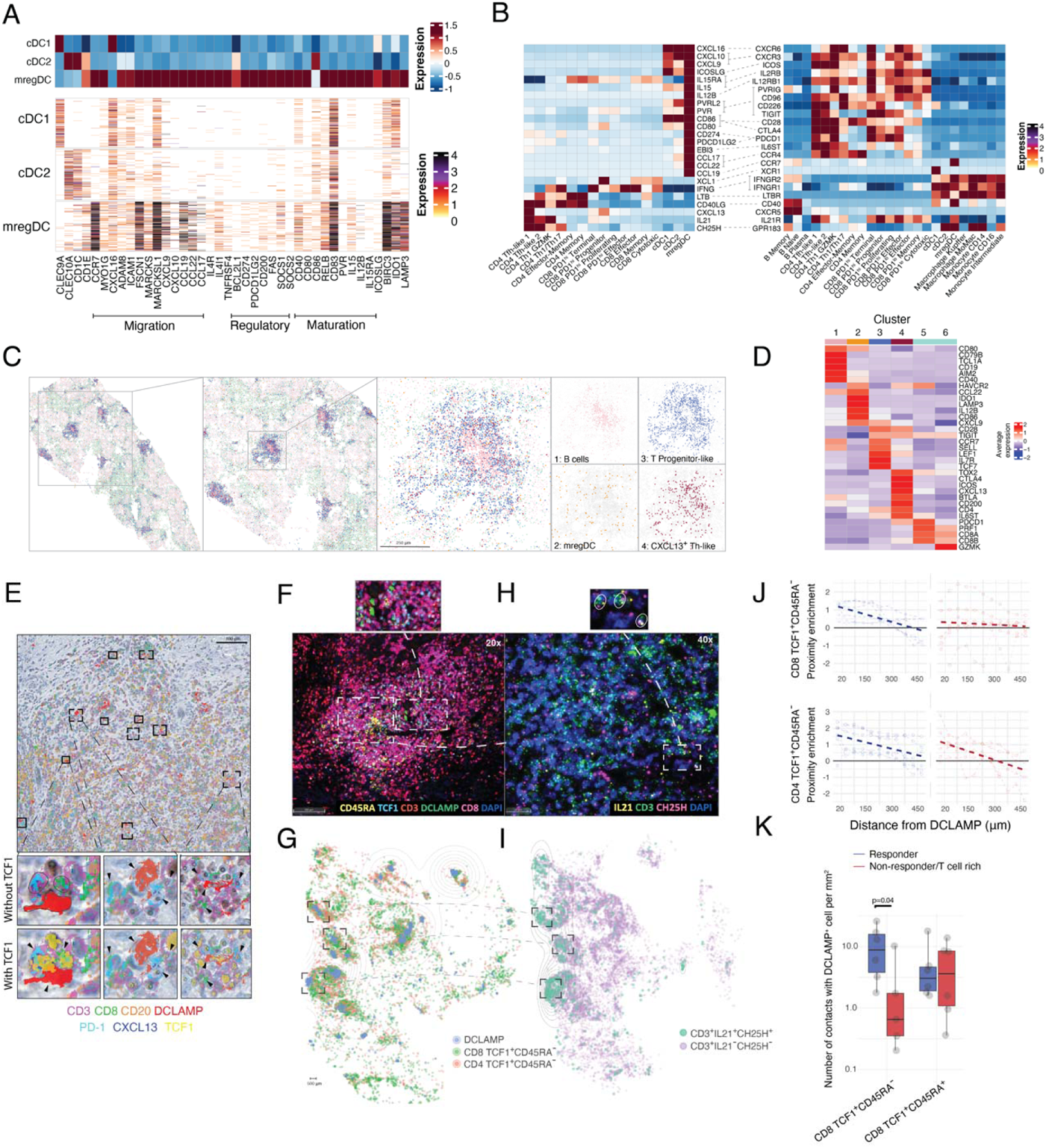
Cellular triads of mregDC, progenitor CD8 and CXCL13 Th cells producing IL-21 and CH25H associate with response to PD-1 blockade. (**A**) Analysis of DC profiles by scRNAseq of cluster-defining genes of DC clusters, showing number of UMI per cell. (**B**) Expression for ligand-receptor pairs of CXCL13 Th and progenitor CD8 T cells across DC, T cell and B cell clusters, showing average expression by scRNAseq. (**C-D**) MERFISH analysis of responder tumor slide (patient 63). (C) Spatial distribution of selected phenotypes at different magnification levels, showing computational rendering of cell localization. (D) Expression of cluster-characteristic genes, showing average expression. (**E-K**) HCC tissue sections of patients treated with PD-1 blockade analyzed by IHC, IF, RNAscope and MICSSS for spatial distribution of T cell subsets and mregDC, post therapy. (**E**) Spatial distribution of mregDC, CD8 and CD4 T cell phenotypes by MICSSS in a representative responder patient 1037. (**F**) Characterization of DC and T cell phenotypes by IF in a representative niche (patient 1017, responder). (**G**) Spatial distribution of mregDC, CD8 and CD4 T cell phenotypes in representative patient (1017, responder), showing computational rendering of IF with density contour annotation for DCLAMP^+^ cells. (**H**) Characterization of CXCL13^+^ Th phenotypes by RNAscope in a representative niche from (**F**). (**I**) Spatial distribution of CD3^+^ IL21^+^ CH25H^+^ T cell phenotypes and CD3^+^ IL21^−^ CH25H^−^/CD3^−^ IL21^+^ CH25H^+^ controls in representative patient from (**G**), showing computational rendering of RNAscope with density contour annotation for CD3^+^IL21^+^CH25H^+^ cells. (**J**) Spatial proximity enrichment of CD8 and CD4 (CD3^+^CD8^−^) T cell phenotypes to DCLAMP^+^ cells by IF at varying distances. (**K**) Quantification of CD8 T cell phenotype contacts (up to 20 μm distance) with DCLAMP^+^ cells.

Using receptor-ligand mapping (**Figure 4B**), we analyzed the expression of candidate molecules that might promote interactions between mregDC, CXCL13^+^ Th, and PD-1^hi^ CD8 T cells. We found that among cDC, mregDC expressed the highest levels of the CD4 T cell ligands (*CCL22*, *CCL17*); the naive and central memory T cell ligand *CCL19*, co-stimulatory molecules (*CD80*, *CD86*, *PVR*, *PVRL2*, *CD40*); cytokines that modulate T cells, such as *IL12B*, which is known to promote Th1 cell differentiation (Garris et al., 2018; Martínez-López et al., 2015; Schurich et al., 2013), and *IL15*, which has been shown to promote CD8 and NK cell survival; and regulatory molecules *CD274* (encoding PD-L1) and *PDCD1LG2* (encoding PD-L2), which we hypothesize may help protect progenitor CD8 T cells from terminal exhaustion (Dähling et al., 2022).

CXCL13^+^ Th cells expressed the highest levels of *IL21*, which sustains the effector function of PD-1^hi^ CD8 T cells (Xin et al., 2015). Interestingly, CXCL13^+^ Th also expressed *CH25H* which may mobilize EBI2-expressing (also known as GPR183) DC, monocyte-derived cells, and progenitor CD8 T cells (Eberhardt et al., 2021; Hudson et al., 2019), and lymphotoxin-β (*LTB*), which plays a key role in the formation of lymphoid structures (Tang et al., 2017) and is known to promote the recruitment of DC and other myeloid populations expressing LTB receptor. CXCL13^+^ Th also expressed high levels of *CD40L*, which promotes the licensing of mregDC and enhance their ability to activate CD8 T cells (Ferris et al., 2020; Laidlaw et al., 2016). In line with prior reports, progenitor PD-1^hi^ CD8 T cells expressed *XCL1*, whose receptor *XCR1* is highly expressed on cDC1 (Brewitz et al., 2017; Crozat et al., 2010; Dorner et al., 2009; Mattiuz et al., 2021).

We then probed spatial interactions between T cells and mregDC using multiplex imaging analysis. We designed a 405 target library for Multiplexed Error-Robust Fluorescence in situ Hybridization (MERFISH, **Table S3**) and performed clustering analysis of 461,533 cells from a full tumor section from a single responder. Strikingly, we found clusters of T cells resembling progenitor-like cells, CXCL13^+^ Th cells, and mregDC that co-localized in discrete cellular niches that were populated by B cells **(Figure 4C-D, S4B)**. Using multiplex immunohistochemistry, we confirmed the accumulation of intratumoral cellular niches that were comprised of DC-LAMP^+^ mregDC, PD-1^+^CXCL13^+^(CD8^−^CD3^+^) CD4 T cells and PD-1^+^ TCF1^+^ (protein encoded by TCF7 gene) progenitor CD8 T cells, along with an abundance of B cells **(Figure 4E, S4C)**. Orthogonal analyses using multiplex immunofluorescence microscopy confirmed that TCF1^+^ CD45RA^−^ progenitors, but not TCF1^+^ CD45RA^+^ naive T cells, accumulated in mregDC niches **(Figure 4F, S4D)**, while TCF1^−^ CD45RA^−^ effector T cells were depleted from these niches **(Figure 4F-G, S4E-F)**. RNAscope of key CXCL13^+^ Th molecules confirmed the localization of *IL21^+^ CH25H^+^*T cells around mregDC niches **(Figure 4H-I)**. Quantitative proximity analysis of full tissue sections indicated that TCF1^+^ CD45RA^−^ CD8 T cells accumulated closer to mregDC in responders compared to non-responders **(Figure 4J, S4G-H)**. Furthermore, the number of TCF1^+^ CD45RA^−^ progenitor-like CD8 T cells localized in close proximity (less than 20µm) of DC-LAMP^+^ mregDC with was higher in responders compared to non-responders **(Figure 4K)**.

Altogether, these results suggest that cellular triads harboring mregDC and CXCL13^+^ Th cells may be critical for promoting local differentiation of PD-1^hi^ progenitor CD8 T cells into potent effector-like CD8 T cells and that these interactions in the TME are essential for the success of PD-1 blockade in HCC.

## DISCUSSION

In this study, we sought to identify molecular correlates of response to PD-1 blockade in T cell-rich HCC tumors, in contrast to T-cell-low and excluded tumors, because despite no apparent defects in T cell priming and recruitment to tumors, a large subset of T cell-rich lesions still failed to respond to ICB.

We observed striking differences between T-cell-rich tumor lesions from responders and non-responders, which we surmised might elucidate the differences in clinical response seen in these patients. Notably, we found that lesions from responders were enriched with CXCL13^+^ Th cells and PD-1^hi^ effector-like CD8 T cells, and that mregDC formed cellular niches with CXCL13^+^ Th cells and PD-1^hi^ progenitor CD8 T cells, which we hypothesize enables their local differentiation into effective anti-tumor CD8 T cells upon PD-1 blockade.

Our results suggest that T cell expansion and activation during PD-1 blockade occurs mostly within the tumor site based on several observations. First, using TCR sequencing and imaging analysis, we found that many T cell clones that expanded after two doses of PD-1 blockade were also present in tumor lesions prior to treatment. Since measurement of TCR diversity prior to treatment was done on minuscule tumor biopsies, it is likely that we may be underestimating the number of pre-existing clones that expand during treatment. Second, using combined scRNAseq/scTCRseq, we show that tumor-expanded CD8 T cell clones consisted of cells at different stages of differentiation, including cells at the progenitor, proliferating and differentiated stages, such as effector-like cells, which dominated in responders or terminally-differentiated cells in non-responders. Collectively, these findings suggest that CD8 T cell differentiation occurs locally within the TME. Finally, while there was a comparable degree of expansion of tumor-enriched clones in the tdLN between responders and non-responders, the prevalence of tumor-enriched clones at the tumor site was much higher in responders, again indicating that local cues regulated the differentiation and expansion of tumor-specific effector T cells, albeit the sampling time following the last treatment with PD-1 blockade could have made the detection of differences in T cell expansion in the tdLN more challenging.

The diversity of PD-1^hi^ CD8 T cells had previously been well described in animal tumor models or during chronic lymphocytic choriomeningitis virus infection, wherein progenitor CD8 T cells proliferate and differentiate into effector-like cells upon PD-1 blockade (Hudson et al., 2019; Im et al., 2016; Miller et al., 2019; Siddiqui et al., 2019). But, efforts to understand their diversity and development in human lesions has been lacking. The high number of cells in our study enabled the identification of the rare CD8 Tpex cells in HCC. These cells expressed PD-1, low levels of naive/memory-associated genes, high levels of *XCL1*/*XCL2*, and genes usually associated with CXCL13^+^ Th cells (e.g. GNG4, BTLA, CD200), consistent with previous studies that identified progenitor CD8 T cells in mouse models (Im et al., 2016). Notably, we found that progenitor CD8 T cells were also present in T-cell-rich lesions from non-responders and displayed clonal overlap with terminally-differentiated cells, suggesting that the lack of response to PD-1 blockade may be due to an inability for progenitor-exhausted CD8 T cells to differentiate into effector CD8 T cells or an inability for effector CD8 T cells to survive locally.

T cell differentiation into effector cells is best instructed by DC, which, in addition to providing TCR engagement, also provide cytokines and costimulatory signals that drive optimal T cell effector function (Cabeza-Cabrerizo et al., 2021; Cancel et al., 2019). We had previously shown that mregDC represent a molecular state induced in both cDC1 and cDC2 upon uptake of cellular debris and that mregDCs are enriched in tumor lesions, likely due to the greater availability of tumor cell-associated antigen cargo (Maier et al., 2020). Here, we show that, similar to our finding in lung cancer (Cohen et al., 2022), mregDC physically interact with CXCL13^+^ Th in discrete niches within T cell rich lesions, and that these hubs also include progenitor CD8 T cells, but not effector cells. We find that progenitor CD8 T cells are enriched in close proximity of mregDC in responders compared to non-responders, suggesting that the direct interactions between progenitor CD8 T cells and mregDC enable effective T cell responses. Indeed, we know that engagement of CD28, which remains expressed on progenitor CD8 T cells, is required for effective CD8 T cell responses upon PD-1 blockade (Duraiswamy et al., 2021; Hui et al., 2017; Kamphorst et al., 2017). Thus, it is likely that upon PD-1 blockade, high expression of CD28 ligands on mregDC, namely CD86 and CD80, enables CD28-dependent activation of progenitor CD8 T cells and their subsequent differentiation into effector CD8 T cells. Additionally, mregDC also express high levels of *IL15* and *IL15RA*, which highlights their potential to be heavily involved in the trans-presentation of IL-15, a cytokine known to promote the survival and maintenance of PD-1^hi^ effector and memory CD8 T cells, respectively (Di Pilato et al., 2021; Mitchell et al., 2010). But, in addition to activation molecules, mregDC also express the highest levels of PD-L1 and PD-L2 among DC, and it is possible that mregDC-mediated PD-L1/PD-1 activation on progenitor CD8 T cells limits their terminal differentiation and permits their maintenance in a progenitor state in these discrete cellular niches. These results align with prior studies showing that tumors enriched in MHC-II^+^ cells and CD8 T cells are less likely to relapse after surgery and that proximity to CD11c^+^ MHC-II^+^ cells may enable CD8 T cells to retain a polyfunctional, progenitor state (Dähling et al., 2022; Duraiswamy et al., 2021; Jansen et al., 2019).

mregDC interactions with CXCL13^+^ Th also likely contribute to the differentiation of progenitor CD8 T cells into effector CD8 T cells in responders. In our survey of tissues, CD4 T cells expanded less than CD8 T cells, making it difficult to study the molecular phenotypic diversity of CD4 T cell clones in greater detail. Nevertheless, TCR sharing between CXCL13^+^ Th cells and cycling T cells, the presence of CXCL13^+^ Th clones prior to treatment and notable similarities between CXCL13^+^ Th cells and the progenitor CD8 T cells (e.g. *PDCD1, TCF7, GNG4, BTLA, CD200*), suggest that CXCL13^+^ Th may serve as an intratumoral CD4 T cell pool that undergoes a proliferative burst upon PD-1 blockade. mregDC may also contribute to the maintenance of a progenitor CD4 T cell pool and to shaping their differentiation into effective CXCL13^+^ Th clones, which we found to be heavily enriched in responders, as opposed to Treg clones, which dominated non-responders. The relatively low number of mregDC in tissues makes it difficult to fully capture their molecular programs in the context of their association with lymphoid aggregates, but enhanced spatial technologies, capable of resolving molecular profiles at the single-cell level, could help decipher DC molecular programs within these niches.

Still, ligand-receptor analysis of scRNAseq datasets helped to reveal potential modes of communication between mregDC and CXCL13^+^ Th cells that likely subsequently shape the differentiation of effector CD8 T cells. For example, enhancement of CD28 signaling induced upon PD-1 blockade could activate CXCL13^+^ Th cells and induce expression of CD40L, which then would engage CD40 on mregDC and permit mregDC licensing and re-activation of CD8 T cells. Furthermore, mregDC also express high levels of EBI3 (a subunit of IL-27 and IL-35) and IL-27 which signals on T cells through GP130/IL6ST promote IL-21 production and proper Th function (Batten et al., 2010; Harker et al., 2013). The production of IL-21 by CXCL13^+^ Th alongside mregDC may represent an important pathway engaged by our proposed triad interaction to support successful responses to PD-1 blockade in cancer, as it has previously been shown for the differentiation of PD-1^+^ effector CD8 T cells during chronic antigen stimulation (Elsaesser et al., 2009; Harker et al., 2013; Yi et al., 2009; Zander et al., 2019). CXCL13^+^ Th cells likely also contribute to the physical positioning of the participants of the cellular triads to promote mregDC/CD4/CD8 T cell interactions. DC-T-B cell interactions in LN are organized according to a discrete chemokine gradient that ensures proper cellular interactions and optimal T cell effector function during antigen presentation (Bajénoff et al., 2007). Oxysterol gradients (generated by CH25H activity) are sensed through EBI2 (*GPR183*) receptor and has been shown to control the positioning of CD4 T cells and cDC2 in peripheral LN (Baptista et al., 2019). Here, we showed that EBI2 is expressed by mregDC, while it was previously reported to be expressed also by progenitor CD8 T cells (Eberhardt et al., 2021; Hudson et al., 2019). *CH25H*, on the other hand, is highly expressed by CXCL13^+^ Th cells, which through oxysterol production, may direct the positioning of mregDC and progenitor CD8 T cells, providing an optimal niche for the differentiation of effective CD8 T cells upon PD-1 blockade. CD4 help for CD8 T cell differentiation is well established and may be particularly important to sustain responses to poorly immunogenic antigens such as tumor antigens.

The production of *CXCL13* and *IL21* by CXCL13^+^ Th cells can promote the recruitment and differentiation of B cell into antibody-producing plasma cells and the presence of tumor-associated B cells and plasma cells have recently been associated with improved outcome in cancer patients (Cabrita et al., 2020; Helmink et al., 2020; Petitprez et al., 2020). Thus, it is possible that upon PD-1 blockade, mregDC promote the activation CXCL13^+^ Th cells and thereby enhance antibody production and contribute to tumor clearance and Fc-mediated enhancement of tumor antigen presentation. Altogether, our results highlight the importance of organized cellular niches within tumors consisting of mregDC and CXCL13^+^ Th cells to promote the survival and differentiation of progenitor CD8 T cells into effective anti-tumor CD8 T cells upon PD-1 blockade.

### Limitations

Diagnosis of HCC is largely based on imaging (such as MRI), so it limits access to pre-treatment tumor tissues and as a result, the number and types of profiling assays that can be done on such scarce tissue are limited. In our study, we were able to perform multiplex imaging and scRNAseq/scTCRseq. But, even so, contemporary spatial technologies are unable to profile rarer populations, such as mregDC, with sufficient depth and resolution, thus limiting our ability to fully capture and characterize their molecular programs. Finally, the lack of experimental animal models that faithfully recapitulate the chronic antigen stimulation that occurs in human neoplastic disease, precludes our ability to probe the presence of these niches in animal models over extended periods of time and the exact role of mregDC and CXCL13^+^ Th cells in tumors.

## ACKNOWLEDGEMENTS

We thank members of the Merad and Brown laboratories at the Marc and Jennifer Lipschultz Precision Immunology Institute at Mount Sinai and the Tisch Cancer Institute for insightful discussions and feedback; and the Mount Sinai Flow Cytometry Core, the Human Immune Monitoring Center, and Biorepository and Pathology CoRE Laboratory of the Icahn School of Medicine at Mount Sinai for support. We thank the patients and their families for participating in the clinical trials.

## AUTHOR CONTRIBUTIONS

AM, PH, AOK, MM conceived the project. AM, MM wrote the manuscript, with contributions from PH, AOK. AM, EK conceptualized computational analyses. AM, PH, AOK, MM designed the molecular profiling or multiplex imaging experiments. AM performed computational molecular and spatial analyses, with additional support from NF, DS, MP, JK, SH, BG, NF, WW, KK, NG, EK. PH, TD, SKS, MB, CC, NM, ASS, JLB, CA, MN, YW, LL performed molecular profiling experiments or design. PH, NF, LT, HS, ST, JLB, ZZ, SW, IF, CP, BK, MD, SG performed multiplex imaging experiments or annotation. CH coordinated the clinical and research teams. YL, EH, SH, RM, JH, KC, NG, RD, AT, SG, MS, TUM, GT provided intellectual input.

## METHODS

### Materials availability

The study did not generate new unique reagents.

### Data and code availability

- Human sequencing data will be available at GEO.
- Any additional information required to reanalyze the data reported in this paper is available from the lead contact upon request.

## EXPERIMENTAL MODEL AND SUBJECT DETAILS

### Human subjects

Peripheral blood as well as samples of lymph node, tumor and non-involved adjacent liver were obtained from specimens of patients undergoing surgical resection at Mount Sinai Hospital (New York, NY) after obtaining informed consent in accordance with a protocol reviewed and approved by the Institutional Review Board at the Icahn School of Medicine at Mount Sinai (IRB Human Subjects Electronic Research Applications 18-00407) and in collaboration with the Biorepository and Department of Pathology. Demographic details and information linking samples to patients are provided in **(Table S1, S4)**.

The single-arm, open-label, phase 2 trial of HCC patients with resectable tumors was registered on ClinicalTrials.gov (NCT03916627, Cohort B). 20 patients were enrolled and received two cycles of cemiplimab before surgical resection as described in (Marron et al., 2022a). We also recruited 9 off-label patients that received 2 to 4 doses of nivolumab prior to surgery. For each specimen, a fragment was formalin-fixed and paraffin embedded (FFPE) for histology, another fragment frozen for RNA/DNA extraction. The remainder of the tissue was directly processed for digestion.

### Bulk TCRseq

Libraries for TCR sequencing were prepared from 100ng total RNA using the SMARTer Human TCR a/b Profiling Kit v2 (Takara Bio). Eighteen cycles were used for each of the 2 semi-nested PCR amplification steps. Sequencing was performed on Illumina NovaSeq (Illumina) by multiplexed paired-read run with 2X251 cycles.

### Whole Exome-seq (library prep and sequencing)

DNAseq libraries were prepared from 50ng gDNA using Twist Library Preparation kit with enzymatic fragmentation and Twist’s Universal Adapter System (Twist Bioscience). Exome capture was performed using the Twist Comprehensive Exome Panel (Twist Bioscience) with IDT’s xGen Hybridization and Wash Kit (Integrated DNA Technologies). Sequencing was performed on Illumina NovaSeq (Illumina) by multiplexed paired-read run with 2X76 cycles.

### Tissue processing

Fresh resected specimens were transported to the lab in R10 media (RPMI with 10% FBS) on ice. For cell dissociation of tumor and non-involved adjacent liver, the samples were perfused with digestion buffer consisting of 0.25mg/mL Collagenase IV (C5138, Sigma-Aldrich) and 0.1mg/mL DNase I (DN25, Sigma-Aldrich), dissolved in R10 and chopped into very small fragments. Tissues were digested for 30min at 37°C with constant shaking at 80rpm and afterwards re-suspended using a 20mL syringe and 16G needle to further break up the tissue, filtered through 100µm followed by 70µm cell strainers. Cell suspensions were spun at 500g for 10min at 4°C. Pellets were re-suspended in 25% Percoll (Cytiva Sweden AB, previously adjusted with 10X PBS) and overlaid on 70% Percoll. Percoll gradient was spun at 400g at room temperature (RT) for 20min with 0 acceleration and 0 break. Middle layer was collected and washed with HBSS (Life Technologies). Pelleted cells were treated with 1X RBC lysis buffer (BioLegend) RT for 2min and washed again with HBSS. The lymph node was chopped in the same digestion media, transferred in a petri dish and incubated for 25min at 37°C without shaking. LN was then dissociated with 18G needle and filtered through 70µm cell strainer. Cell suspension was spun at 350g for 5min at 4°C and pellet was incubated in RBC lysis for 2min.

Cells from all samples were resuspended in appropriate buffer for counting and to be used in subsequent assays.

Peripheral blood was obtained the day of surgery for processing and analysis of the peripheral blood mononuclear phagocytes (PBMC). The sample was diluted in PBS and added carefully on top of Ficoll (Ficoll-Paque PLUS, GE Healthcare). Ficoll gradient was spun at 1200g at room temperature (RT) for 10min with 0 acceleration and 0 break. The layer containing the PBMC was collected and washed in PBS. After centrifugation, the pellet was resuspended in appropriate buffer for counting and to be used in subsequent assays.

### CITEseq hashing and staining

Cells were counted using the Nexcelom Cellaca. Aliquots of 400,000 cells from each sample were centrifuged at 350g for 5 minutes at 4°C. The supernatant was discarded and each cell pellet was resuspended in a unique hashtag antibody solution and incubated on ice for 20 minutes. Hashtag antibodies were conjugated as per New York Genome Center hashing protocol. Stained cells were washed three times in 1mL wash buffer (PBS + 0.5% BSA) by 4°C centrifugation at 350g to remove unbound antibodies. Washed cells were resuspended in 150µl wash buffer and counted using a Nexcelom Cellaca. Hashed samples were pooled and centrifuged at 350g for 5 minutes at 4°C. Supernatant was removed and pellet was resuspended in 100µl of antibody cocktail **(Table S5)** and incubated at 4°C for 30 minutes. Stained cells were washed three times in wash buffer by 4°C centrifugation at 350g to remove unbound antibodies. The washed cells were resuspended in wash buffer to achieve a target concentration of 4 million cells/mL.

### CITEseq and scRNAseq processing

For scRNAseq (directly load), cells were counted and loaded on the 10x Genomics 3’v2, 3’v3 or NextGem 5’v1 assay as per the manufacturer’s protocol with a targeted cell recovery of 10,000 cell per lane. For CITEseq, sample pool was counted and loaded on the 10x Genomics NextGem 5’v1 assay as per the manufacturer’s protocol with a targeted cell recovery of 25,000 cells per lane. Gene expression and Feature Barcode libraries were made as per the 10x Genomics demonstrated protocol. Hashtag oligonucleotides (HTO) were enriched during cDNA amplification with the addition of 3pmol of HTO Additive primer (5’GTGACTGGAGTTCAGACGTGTGCTC). This PCR product was isolated from the mRNA-derived cDNA via SPRISelect size selection, and libraries were made as per the New York Genome Center Hashing protocol. All libraries were quantified via Agilent 2100 hsDNA Bioanalyzer and KAPA library quantification kit (Roche). Gene expression libraries were sequenced at a targeted depth of 25,000 reads per cells. ADT libraries were sequenced at a targeted read depth of 10,000 reads per cell. HTO libraries were sequenced at a targeted read depth of 1,000 reads per cell. Paired-end sequencing was performed on Illumina NovaSeq 6000 for RNAseq and CITEseq libraries (for 3’ libraries, used Read 1 28 bp for UMI and cell barcode, Read 2 80-bp for transcript read, with 8-bp i7 and 0-bp i5 reads; for 5’ libraries, used Read 1 26 bp for UMI and cell barcode, Read 2 80-bp for transcript read, with 8-bp i7 and 0-bp i5 reads). V(D)J libraries were also sequenced on Illumina NovaSeq 6000 (Read 1 150-bp, 8-bp i7, 0-bp i5, Read 2 150-bp). Hashtag libraries were sequenced on Illumina NextSeq500.

### Memory T cell enrichment

Cryopreserved PBMC and dissociated tdLN cells were thawed and rested for overnight in GMP-grade Dendritic Cell Media (Cellgenix 20801-0500) containing 5% human AB serum (Sigma H3667) and 1% Penicillin-Streptomycin-Glutamine (Gibco 10378016). Following the incubation of cells with Human Seroblock (Bio-Rad BUF-070B), dissociated LN and PBMC samples were individually stained with unique TotalSeq-C anti-human Hashtag antibodies (BioLegend) as per manufacturer’s instructions. After washing 3 times with Stain Buffer containing BSA (BD Bioscieces 554657), hashed PBMC samples were pooled at equal cell numbers. LN samples and pooled PBMC samples were individually stained with a panel of PE-conjugated oligo-barcoded dCODE Dextramers (10x, Immudex) as per manufacturer’s instructions except that 0.5 μL of each dCODE Dextramer was used to stain 1 ~ 2×10^6^ cells. Dextramer stained cells were washed and blocked with Human Seroblock. Cells were then stained with a TotalSeq-C Custom CITEseq Antibody Cocktail (custom-designed, BioLegend) and the following antibodies: CD45 (Clone HI30, BIoLegend), CD3 (Clone SK7, BD), CCR7 (Clone G043H7, BioLegend), CD45RO (Clone UCHL1, BD), CD95 (Clone DX2, BD), CD19 (Clone SJ25C1, Invitrogen), CD56 (Clone NCAM16.2, BD). T cells were identified as CD45^+^ CD3^+^ CD19^−^ CD56^−^ cells. Memory T cells were sorted using the following 3 sort gate parameters and then pooled for single-cell sequencing: CD45^+^ CD3^+^ CD45RO^+^, CD45^+^ CD3^+^ CD45RO^−^ CCR7^−^, and CD45^+^ CD3^+^ CD45RO^−^ CCR7^+^ CD95^+^. To detect the binding of Cemiplimab to PD-1, an oligonucleotide-barcoded anti-human IgG4 (Clone HP6025, SouthernBiotech) was included in the CITEseq stain panel. Barcoded anti-human IgG4 was generated as previously described (van Buggenum JAGL 2016; Stoeckius M 2018) using an oligo with 5’-amine modification (IDT). DAPI was added to cells prior to FACS on the Sony MA900 instrument. Memory T cells from PBMC and lymph nodes were sorted separately. Aliquots of sorted cells were removed and stored in TRIzol for bulk RNA sequencing. Memory T cells enriched from lymph nodes and PBMC were pooled proportionally for downstream processing. 15,000 to 30,000 cells were loaded onto a Chromium Single Cell 5’ Chip and processed through the Chromium Controller (10x Genomics) for single cell sequencing.

### Gene selection for MERFISH

To identify transcriptionally distinct cell population with MERFISH, we designed a panel of informative genes. Selection of these genes were based on two categories. Category one genes were manually picked to serve as markers for different immune cells including macrophages, T cells, B cells, and dendritic cells, and to serve as functional readout of those cell types including T-cell exhaustion, proliferation, signaling etc. Category two genes were chosen based on previously generated single cell sequencing data and identification of differentially expressed genes of interest. We evaluated this gene panel using the MERSCOPE Gene Panel Design Portal available at Vizgen (portal.vizgen.com) to ensure that each gene is sufficiently long to allow enough encoding probes to bind, and that the entire gene panel meets the abundance threshold to avoid optical crowding for MERSCOPE imaging. This ended in a final panel of 405 genes **(Table S3)**. To serve as a control for unspecific binding of probes, we included 50 blank barcodes.

### Tissue preparation for MERFISH

#### Tissue Sectioning and Permeabilization

Samples from HCC patients were snap frozen and preserved in optimal cutting temperature (OCT) compound and stored at −80°C until sectioning. Frozen tumor samples were sectioned at −20°C on a cryostat (Microm HM525, Thermo Scientific) at 10µm-thick and placed on MERSCOPE Slide (Vizgen 20400001). After fixation with 4% paraformaldehyde in PBS for 15min, tissue slices were washed three times with 5mL PBS and placed in 5mL of 70% ethanol to allow for tissue permeabilization at 4°C overnight.

### Cell Boundary and Antibody Stain

Following overnight permeabilization, patient tissue slices were photobleached using the MERSCOPE Photobleacher (Vizgen 1010003) for 4 hours to remove background fluorescence. Samples were stained for cell boundary using Vizgen’s Cell Boundary Kit (10400009) following Vizgen’s User Guide for fresh and frozen tissue sample preparation (https://vizgen.com/resources/user-guides/). Briefly, the tissue slices were blocked with Blocking Solution (PN 20300012) supplemented with RNase inhibitor (NEB, M0314L) at 1:20 dilution for one hour, washed with PBS and then incubated with primary antibody from the Cell Boundary Primary Staining Mix (PN 20300010) and CD68 antibody (Agilent Dako) at 1:100 dilution for 1 hour with the supplement of RNase inhibitor (NEB, M0314L) at 1:20 dilution. Afterwards, we further incubated with ready-to-use CD3 antibody (Ventana) supplemented with RNase inhibitor (NEB, M0314L) at 1:20 dilution for another 1 hour. After washing with PBS three times, the samples were stained with oligo conjugated secondary antibodies supplemented with RNase inhibitor (NEB, M0314L) at 1:20 dilution for 1 hour, post-fixed with 4% paraformaldehyde in PBS for 15 minutes, washed with PBS and next prepared for MERFISH encoding probe hybridization.

### MERFISH Encoding Probe Hybridization

Samples were washed with 5mL Vizgen Sample Prep Wash Bugger (Vizgen 20300001) for 5 minutes and incubated at 37°C in 5mL Formamide Wash Buffer (Vizgen 20300002) for 30 minutes. After aspirating the buffer, we applied 50µL of custom designed MERSCOPE Gene Panel Mix (Vizgen 20300008) to the tissue, covered with parafilm to prevent evaporation and placed in a 37°C incubator for 36-48 hours. Following incubation, tissues were washed twice with 5mL Formamide Wash Buffer at 47°C for 30 minutes and finally washed with 5mL Sample Prep Wash Buffer for 2 minutes.

### Gel Embedding and Tissue Clearing

Samples were embedded into a gel made from 100µL of gel embedding solution. Gel embedding solution was made with 5mL of Gel Embedding Premix (Vizgen 20300004), 25µL of 10% ammonium persulfate (Sigma, 09913-100G) and 2.5µL of TEMED (N,N,N’,N’-tetramethylethylenediamine) (Sigma, T7024-25ML). 20mm Gel Coverslips (Vizgen 20400003) were prepared with RNAseZap, 70% ethanol and covered with 100µL Gel Slick (VWR, 12001-812). Samples were incubated with 5mL of the gel solution for 1 minute and following removal of the solution, 100µL of gel solution was added on top of the sample and sandwiched beneath the Gel Coverslip. Excess gel solution was aspirated, and the samples incubated at room temperature for 1.5h to allow the gel solution to polymerize. After removing the Gel Coverslip, the samples were incubated with Clearing Solution consisting of 50µL of Protease K (NEB, P8107S) and 5mL of Clearing Premix (Vizgen 20300003) at 47°C overnight first and then at 37°C for one additional night. A fully detailed, step-by-step instruction on the MERFISH sample prep the full protocol is available at https://vizgen.com/resources/fresh-and-fixed-frozen-tissue-sample-preparation.

### Sample Imaging

Clearing solution was removed, and samples were washed for 10 minutes with Sample Prep Wash Buffer. Samples with incubated with 3mL of DAPI and Poly T Reagent (Vizgen 20300021) for 15 minutes at room temperature, washed for 10 minutes with 5mL of Formamide Wash Buffer and transferred to 5mL of Sample Prep Wash Buffer. The imaging buffer was prepared by adding the Imaging Buffer Activator (Vizgen 20300015) and RNase inhibitor to the imaging buffer. The imaging reagents and processed samples were loaded to the MERSCOPE system (Vizgen 10000001). Following a low-resolution DAPI mosaic at 10x magnification the regions of interest were selected for high-resolution imaging at 60x. Full Instrumentation protocol is available at https://vizgen.com/resources/merscope-instrument/. Data were generated and use for cell segmentation and analysis.

### MICSSS

FFPE tissue sections (4µm) were stained using the multiplexed immunohistochemical consecutive staining on a single slide (MICSSS) protocol as previously described (Remark et al., 2016). Briefly, slides were baked at 50°C overnight, deparaffinized in xylene and rehydrated in decreasing concentration of ethanol (100%, 90%, 70%, 50% and dH2O). Sample slides were incubated in pH6 or pH9 buffers at 95°C for 30min for antigen retrieval, then in 3% hydrogen peroxide for 15min and in serum-free protein block solution (Dako) for 30min. Primary antibody staining was performed using the optimized dilution during 1h at room temperature (RT) or at 4°C overnight followed by signal amplification using associated secondary antibody conjugated to horseradish peroxidase (HRP) during 30min. Chromogenic revelation was performed using AEC (Vector). Tissue sections were counterstained with hematoxylin, mounted with a glycerol-based mounting medium and finally scanned to obtain digital images (Aperio AT2, Leica). After scanning, slide coverslips were removed in hot water (~50°C) and tissue sections were bleached and stained again as previously described (Remark et al., 2016). Primary antibodies are presented in **(Table S6)**.

### Multiplex IHC

A fully automated Multiplex Immunohistochemistry assay was performed on the Ventana Discovery ULTRA platform (Ventana Medical Systems, Tucson, AZ), and as previously described (Marron et al., 2022a). The assay was optimized for HCC. Optimal concentrations of each antibody were determined, and they were applied in the following sequence and detected with the indicated fluorophore **(Table S7)**. Following staining, the tissue was counter-stained and cover slipped with Invitrogen ProLong Gold Antifade Mountant with NucBlue. Whole slide imaging was performed on the Zeiss Axioscan which was equipped with a Colibri light source and appropriate filters for visualizing these specific fluorophores.

### BaseScope in situ TCR detection

We performed the Advanced Cell Diagnostics (Biotechne Brand, Minneapolis, MN) BaseScope VS assay on the Roche/Ventana Discovery ULTRA automated staining instrument (Ventana Medical Systems, Tucson, AZ). The BaseScope assay uses a novel and proprietary method of in situ hybridization to visualize single RNA molecules. Probes are specifically designed to bind as pairs and for BaseScope, a single ZZ pair is amplified using multiple steps. This is followed by hybridization to alkaline phosphatase-labeled probes and detection using a red chromogenic substrate. Fresh 4µm tissue sections were prepared on Fisherbrand Superforst PLUS slides (Cat. No. 12-550-15) and the fully automated BaseScope protocol was followed. Samples were pre-treated with Target Retrieval solution and a protease step to expose the RNA and to allow the RNA-specific probes to hybridize to their target RNA. Negative and positive control probes were included to assess the quality of the RNA (Negative Control Probe: DapB-1ZZ and Positive Control Probe: Human-PPIB-1ZZ). Probes were custom designed for the individual patient based on four chosen CD4 and CD8 TCR sequences and assayed as a CD4 and CD8 cocktail. On completion of the 12 hours staining run, slides were counterstained with Novacastra Hematoxylin (Leica Microsystems Inc.), dried in a 60°C oven for 30 minutes and after a brief dip in Xylene, were mounted with EcoMount (Biocare). All slides were scanned on the Leica Aperio AT2 scanner at 40x (Leica Biosystems, Deer Park, IL) and images were analyzed using HALO Indica Labs modules (IndicaLabs).

### Multiplex fluorescent RNAscope

The Advanced Cell Diagnostics (Biotechne Brand) RNAscope LS Multiplex Fluorescent Assay was performed on the Leica BOND RX automated staining system (Leica Biosystems). The Multiplex Fluorescent assay involves three independent signal amplification systems each using a different fluorophore (Akoya Biosciences) to visualize the target RNA (TSA Plus Fluorescein, TSA Plus Cyanine 3 and TSA Plus Cyanine 5). Fresh 4µm tissue sections were prepared on Fisherbrand Superforst PLUS slides and the fully automated Multiplex RNAscope protocol was performed. Samples were pre-treated with Target Retrieval solution and a protease step to expose the RNA and to allow the RNA-specific probes to hybridize to their target RNA. Control probes were run on all samples to assess the quality of the RNA (RNAscope LS Multiplex Negative Control probe and Positive control Probe). Target-specific probes were pooled and diluted appropriately to stain for *CD3E, IL21* and *CH25H* **(Table S8).** Tissues were counterstained with DAPI to allow for visualization of the nuclei.On completion of the 14 hour run, slides were cover slipped with ProLong Gold Antifade Medium (Thermo Fisher). Slides were scanned on the Zeiss Axioscan.Z1 scanner at 20x magnification with Z-stacking (Zeiss). Quantitative image analysis was preformed using HALO Indica Labs modules (IndicaLabs).

## QUANTIFICATION AND STATISTICAL ANALYSIS

### Analysis of sequencing data

For scRNAseq libraries, Cell Ranger Single-Cell Software Suite (10X Genomics, v2.2.0) was used to perform sample demultiplexing, alignment, filtering, and UMI counting. The human GRCh38 genome assembly and RefSeq gene model for human were used for the alignment. For V(D)J libraries, Cell Ranger Single-Cell Software Suite (10X Genomics, v2.2.0) was used to perform sample de-multiplexing, de novo assembly of read pairs into contigs, align and annotate contigs against all of the germline segment V(D)J reference sequences from human IMGT, label and locate CDR3 regions, group clones.

### Unsupervised batch-aware clustering analysis

Immune cells from tumor and adjacent tissue samples were filtered for cell barcodes recording > 500 UMI, with < 25% mitochondrial gene expression, and with less than defined thresholds of expression for genes associated with red blood cells and epithelial cells. Cells were clustered using an unsupervised batch-aware method we recently described (Leader et al., 2021) with minor adjustments. This EM-like algorithm, which was also based on earlier studies (Jaitin et al., 2014; Paul et al., 2015), iteratively updates both cluster assignments and sample-wise noise estimates until it converges, using a multinomial mixture model capturing the transcriptional profiles of the different cell-states and sample specific fractions of background noise. As opposed to other standard clustering approaches, this probabilistic method explicitly distinguishes between the measurements of transcripts from individual cells from generative models of expression built using cluster averages and noise approximations. The use of such a model facilitates both the clustering of cells from distinct batches as well as the classification of cells from additional data that were not used in the initial clustering. We clustered tumor and adjacent samples from 16 patients (73 and 61 samples, respectively) sequenced prior to Feb 28^th^, 2021, and then mapped additional samples onto the final model as described below.

The model definitions and estimation of model parameters were as described in (Martin et al., 2019). The multinomial mixture model of gene expression assumes that the expression of cells from each cluster can be modelled as a multinomial distribution, with each gene having a specific probability of sampling, and that these probabilities are uniform for all cells in a cluster. Here, we further modified the probabilities using a batch-specific term in order to account for batch noise that was observed in the data and was modelled as the average expression of the batch.

We also used here the pseudo expectation-maximization (EM) algorithm (Martin et al., 2019) to infer the model parameters with minor modifications: (1) training and testing set size were 4000 and 2000 respectively and (2) the best clustering initiation was selected from 5000 kmeans+ runs. Genes with high variability between patients were not used in the clustering. Those genes consisted of mitochondrial and stress genes, metallothionein genes, immunoglobulin variable chain genes, HLA class I and II genes and 3 specific genes with variable/noisy expression: *MALAT1*, *JCHAIN and XIST* **(Table S9)**. Ribosomal genes were excluded only from the k-means clustering in step 2.D, as described in (Martin et al., 2019).

### CITEseq analysis

Raw counts of CITEseq data were normalized by DSB (denoised and scaled by background) normalization method (Mulè et al., 2022). DSB normalization uses background droplets to evaluate protein background noise to correct values in cells and quantify protein counts above those background levels in individual cells. DSB also uses isotype control and each cell’s specific background level to remove technical cell-to-cell variation. Demultiplexed cells with designation as negative were used as background droplets while demultiplexed singlets were used as positive cells. 6 Isotype controls from the CITEseq panel were used as DSB isotype control input in the modeled DSB technical component. DSB normalization was then performed with aforementioned raw counts matrix from positive cells and background droplets as well as isotype controls.

### Statistical testing of differential abundances

Abundances were calculated as frequency relative to the discussed compartment. Two-tailed T tests were used for significance assessment of differential abundance. Significance was determined as p<0.05. Multiple hypotheses correction (Benjamini & Hochberg) was applied when applicable.

### Analysis of public datasets

UMI data was downloaded from GEO and average expression per CXCL13^+^ Th1/Tfh CD4 and non-CXCL13^+^ Th1/Tfh CD4 clusters (as described in (Zheng et al., 2021)) was calculated for each CD4 T cell marker.

### Gene-modules analyses

As previously described (Leader et al., 2021; Martin et al., 2019), cells were down-sampled to 2000 UMI prior to selecting a set of variable genes and gene-gene correlation matrix was computed for each sample for CD8 and separately CD4 T cells. Correlation matrices were averaged via Fisher Z-transformation. The inverse transformation then resulted in the best-estimate correlation coefficients of gene-gene interactions across the dataset. Genes were clustered into modules using complete linkage hierarchical clustering over correlation distance.

### scTCRseq analysis

Single T cells were grouped by clonotype according to their precise combination of alpha and beta chains present (uniquely defined by CDR3 sequence and V, D, and J gene usage), with the following exceptions to filter for high quality singlets:

1. Cells with contigs encoding > 3 productive α and β chains were excluded as multiplets.
2. Cells with contigs encoding >3 productive α and β chains that completely overlapped with observed cells within the multiplets were also excluded as multiplets.
3. Remaining cells with 3 unique α and β chains that could be uniquely associated with similar cells displaying 2 unique α and β chains were assumed to be clonally related, whereas cells that could be similarly associated with multiple distinct sets of cells expressing 2 unique α and β chains were excluded as doublets.
4. Cells in which a single TCR chain was observed were assumed to be clonally related to any cells with 2 unique α and β chains to which they uniquely associated.
5. Remaining cells in which a single TCR chain was observed were excluded if they matched ambiguously to multiple cells with 2- or 3-chains.

Clones with 2 or more cells were termed expanded, in contrast to those with only a single cell which were termed singlets. Clones expanded in only a single tissue (tumor or adjacent) which displayed 1.5 or more fold-change of clone abundance relative to the other tissue, were deemed tumor- or adjacent-enriched, respectively. The statistical significance of tissue enrichment per clone was determined using a permutation test, whereby the tumor and adjacent tissue origin was shuffled for 1000 repetitions, and the relative abundance of each clone was compared across tumor and adjacent tissues. In contrast to CD8 T cells which are clonally expanded to a large degree and where we had sufficient power to identify statistically significant tumor-enrichment, the relatively limited clonal expansion of CD4 T cells restricted such filtering and therefore we included all tumor-enriched clones in the analysis regardless of significance. Top tumor-enriched clones for TCR sharing visualization, BaseScope imaging, bulk TCRseq and LN scTCRseq analyses were selected based on clone size, and further filtered to include only those containing at least 10% of CXCL13^+^ Th, or separately, PD-1^hi^ Effector CD8 T cells. The selection for BaseScope imaging was further manually curated for clones dominated by the CD4 and CD8 population of interest. Tversky index and Gini inequality index were calculated over the combined patient data including all tumor-enriched clones.

### Ligand-receptor analysis

Ligands or receptors for CXCL13^+^ Th, PD-1^hi^ CD8 T cell and mregDC marker genes were identified via CellPhoneDB, which aggregates cellular communication axes across UniProt, Ensembl, PDB, the IMEx consortium, IUPHAR. The average expression of matching receptors or ligands was visualized for cell types of interest, including B cells, CD8 and CD4 T cells, and DC.

### Somatic mutation calling and tumor mutation burden calculation

Somatic mutations were called using the Sentieon somatic FASTQ to VCF (version 3.2.0) applet with mark duplicates option, TNscope algorithm selected for mutation calling, extra BWA option –K 10000000, and extra TNscope options --clip_by_minbq 1 --max_error_per_read 3 --min_init_tumor_lod 2.0 --min_base_qual 10 -- min_base_qual_asm 10 --min_tumor_allele_frac 0.00005. Machine learning (ML) model SentieonTNscopeModel_GiAB_HighAF_LowFP-201711.05.model was utilized during mutation calling.

Called somatic mutations that had ML model probability of less than 0.81, were reported in dbSNP151 (common), had variant allele frequency in tumor less than 0.07, did not have at least 40 read coverage, did not have at least four variant supporting reads in tumor, or had more than one variant supporting read in germline were filtered out.

After mutation filtering, variant consequence was annotated using SnpEff (version 4.3t) and tumor mutation burden was calculated separately for all mutation types combined, indels only, nonsense mutations only, and missense mutations only.

### MERFISH Cell Segmentation and Analysis

Cell segmentation was performed by first identifying nuclear seeds using the convex object identification library Stardist, second identifying cell boundary peaks along radial paths from nuclear seeds, third defining polygons connecting cell boundary peaks, and fourth connecting overlapping polygons from different Z slices to construct 3D segmented cells. Single-cell analysis was performed using the Scanpy library. MERFISH single-cell gene expression data was first filtered to remove cells with low count or low number of unique genes expressed (5 genes and 10 counts respectively). Cells were then normalized to have equal counts (set to default of median of total counts) and gene expression counts were log transformed and scaled to unit variance. PCA, UMAP embedding (10 neighbors, 0.1 min distance, and spread 3.0), and Leiden clustering (resolution of 1.5) were performed, and Leiden clusters were manually labeled using their average expression.

### Multiplex IHC

Quantitative image analysis was preformed using HALO Indica Labs Hyperplex module (IndicaLabs). Number of positive cells for each immune subset and density in the entire tumor area was measured. In the absence of a reliable liver tumor marker, we performed the analysis across tumor regions annotated by expert pathologists.

### Imaging proximity analysis

Separately for CD4 and CD8 T cell phenotypes, a spatial K nearest neighbor (KNN) network was constructed using the Giotto R package, with a maximum distance limit of 500μm. Then the proximity enrichment for phenotype X was assessed for each tier of increasing distance (0-20, 0-150μm etc, as shown in figure) as the ratio of observed frequency of X neighboring DC-LAMP to the expected frequency based on X abundance in the general population regardless of DC-LAMP proximity.

---

**Figure S1.**
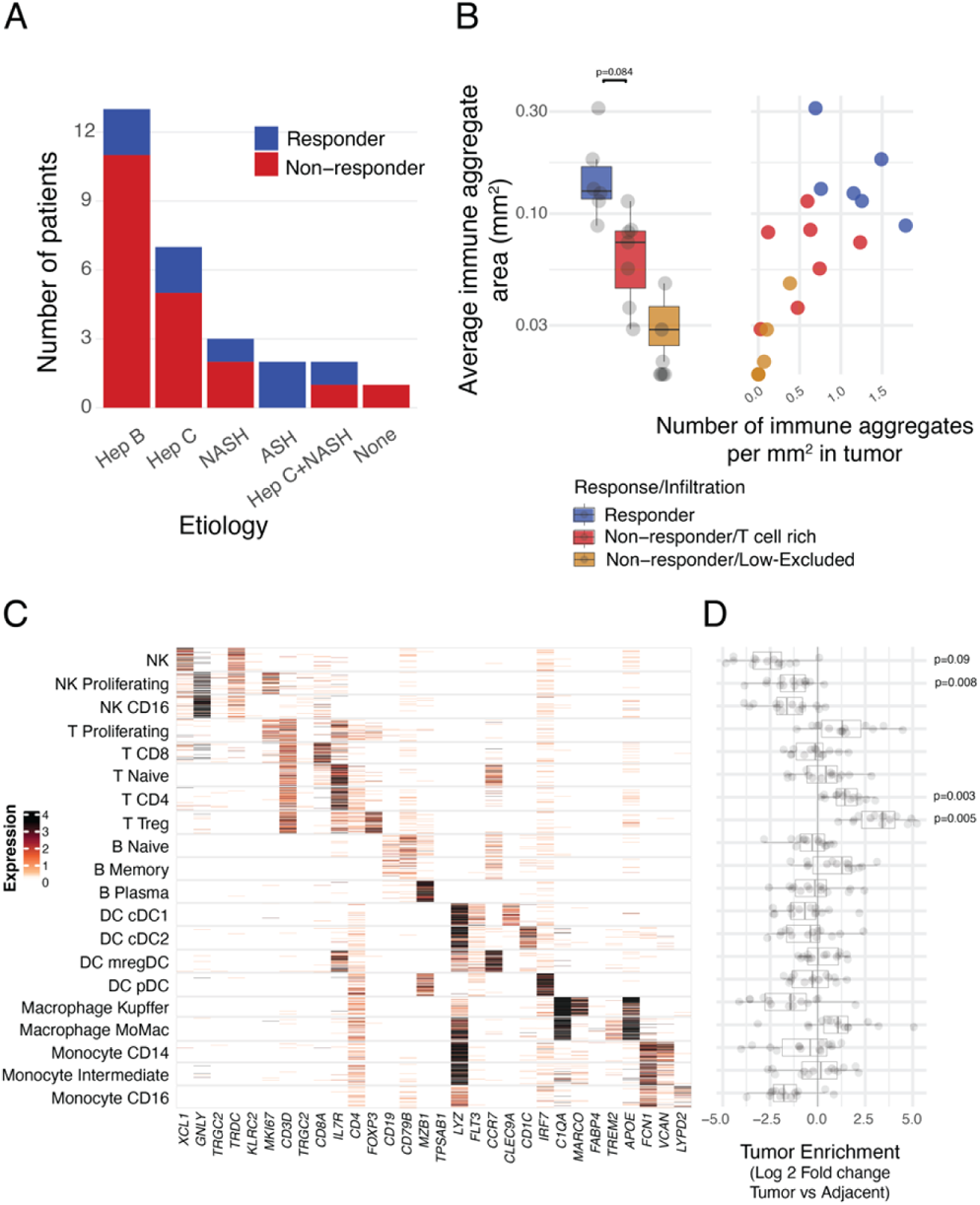
Characterization of T cell rich HCC lesions in response to PD-1 blockade. (**A**) Distribution of responders and non-responders across HCC etiologies (Hep B: Hepatitis B; Hep C: Hepatitis C; NASH: Non-alcoholic steatohepatitis; ASH: Alcoholic steatohepatitis). (**B**) Quantification of immune aggregate areas and numbers stratified by response and immune infiltration pattern. (**C**) Expression of cluster-defining genes by scRNAseq of key immune populations, showing number of UMI per cell. (**D**) Differences of cluster frequencies between tumor and adjacent tissue.

**Figure S2.**
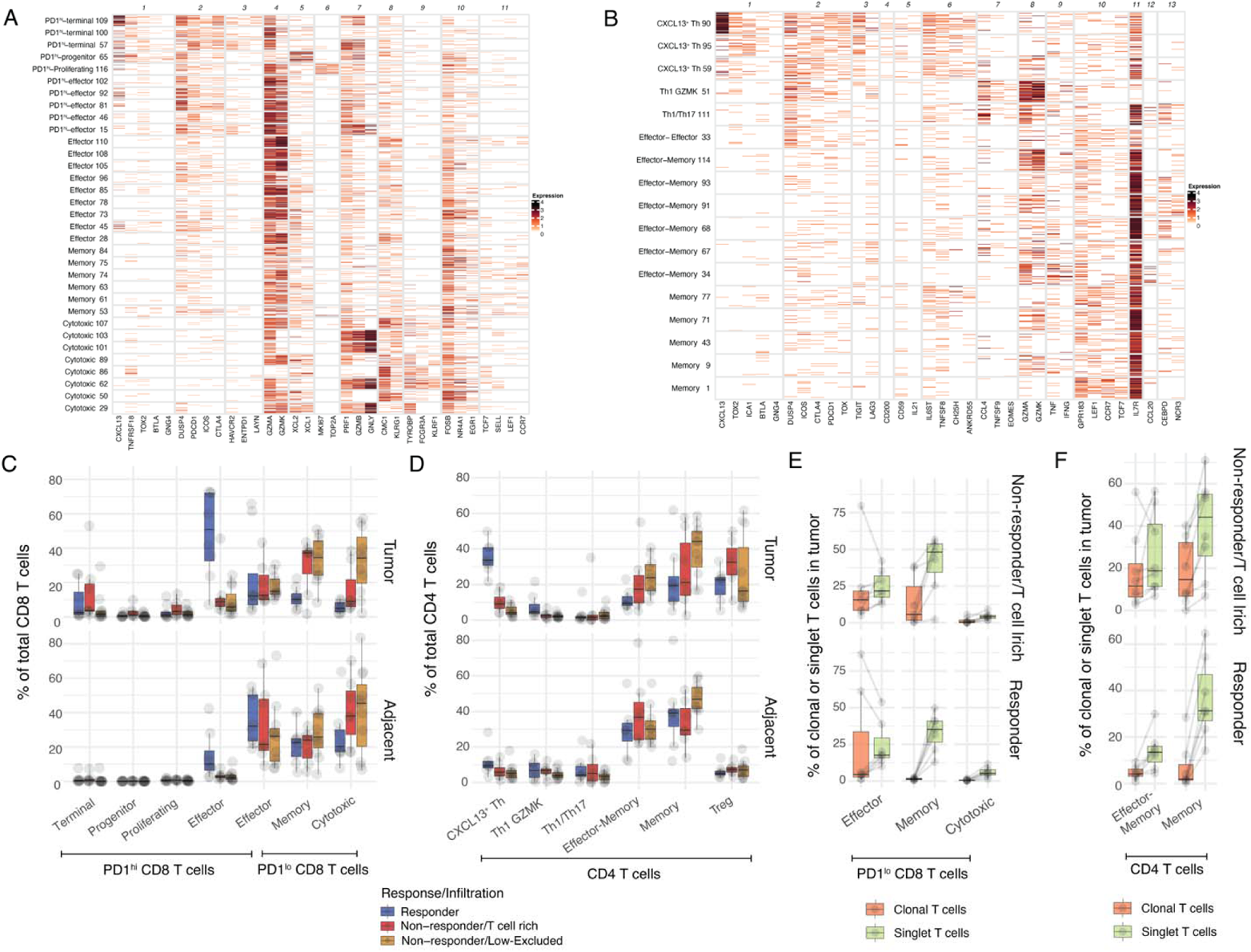
Molecular profiling of expanded CD8 and CD4 T cell clones associated with response to PD-1 blockade. (**A-B**) Expression of cluster-defining gene modules by scRNAseq for (A) CD8 (B) CD4 T cell clusters showing number of UMI per cell. (**C-D**) Cluster frequencies among (**C**) CD8 and (**D**) CD4 T cells in tumor and adjacent tissue, stratified by response and immune infiltration pattern. (**E**) Cluster frequencies among PD-1^lo^ CD8 T cells in tumor among tumor-enriched clonal T cells and tumor singlet T cells, stratified by response and immune infiltration pattern. (**F**) Cluster frequencies among CD4 T cells in tumor among tumor-enriched clonal T cells and tumor singlet T cells, stratified by response and immune infiltration pattern.

**Figure S3.**
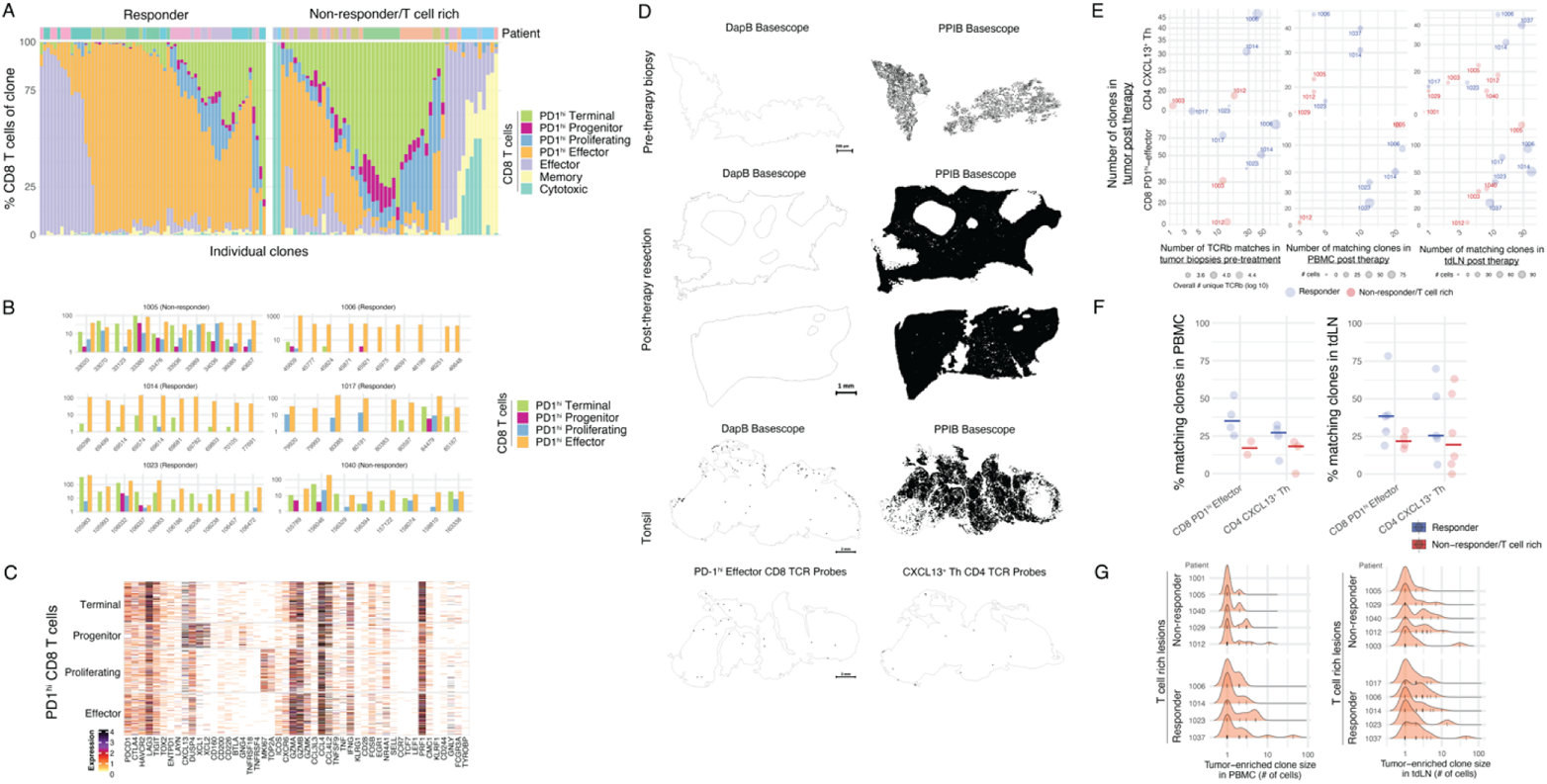
Phenotypic distribution among CD8 T cell clones and clonality assessment in PBMC and tdLN. (**A-C**) Phenotypic analysis of clonotype sharing by scTCRseq. (**A**) Phenotypic distribution of all CD8 T cell clusters in individual tumor-enriched clones (top 10 per patient), separately for responders and T cell rich non-responders. (**B**) Highlight of PD-1^hi^ CD8 phenotypic distribution for selected CD8 clones across remaining patients not shown in Figure 3B. (**C**) Expression of PD-1^hi^ CD8 cluster-defining genes by scRNAseq for CD8 T cells from the selected clones of Figure 3B for patient 1037 (responder), showing number of UMI per cell. (**D**) BaseScope TCR imaging DapB and PPIB controls across biopsy, resection and tonsil samples. (**E-G**) scTCRseq of tumor, PBMC and tdLN from time of resection. (**E**) Number of post-treatment tumor-enriched clones present in pre-treatment tumor lesions by bulk TCRseq, and PBMC and tdLN by scTCRseq, across responders and T cell rich non-responders. (**F**) Percent of post-treatment tumor-enriched clone present in scTCRseq of PBMC and tdLN, across responders and T cell rich non-responders. (**G**) Histograms of clone size (number of cells per clone) distribution per patient, stratified by response and immune infiltration pattern, separately in PBMC and tdLN. Ticks represent individual clones.

**Figure S4.**
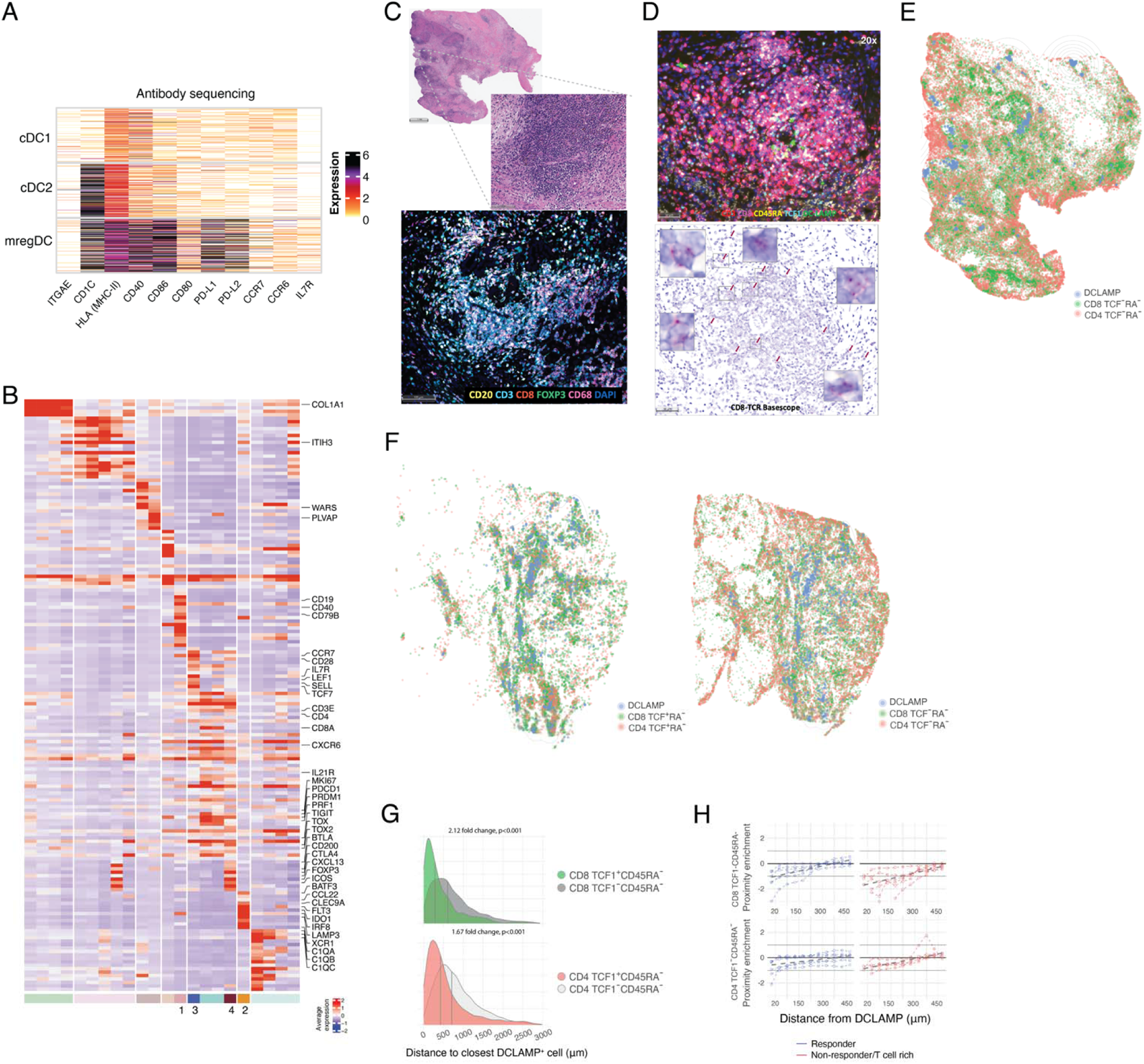
Spatial localization of CXCL13 Th, progenitor CD8 T cells and mregDC. (**A-H**) HCC tissue sections of patients treated with PD-1 blockade analyzed by MERFISH, IHC, IF and BaseScope for spatial distribution of T cell subsets and mregDC, post therapy. (**A**) Expression of DC cluster-defining proteins by CITEseq antibody sequencing, showing number of UMI per cell. (**B**) Expression of cluster-characteristic genes from MERFISH analysis, showing average expression. (**C**) IHC analysis of macrophages, T and B cells in representative responder (patient 1017). (**D**) IHC (top) and BaseScope CD8 TCR localization (bottom) in a representative immune aggregate in responder (patient 1006). (**E-F**) Spatial distribution of mregDC, CD8 and CD4 phenotypes in representative responder patients (**E**) 1017 and (**F**) 1041, showing computational rendering of IF with density contour annotation for DCLAMP^+^ cells. (**G**) Distribution proximity to closest CD8 and CD4 T cell phenotypes from DCLAMP^+^ cells, showing histograms of individual cells from patient 1017, vertical gray bars represent the median. (**H**) Spatial proximity enrichment of CD8 and CD4 (CD3^+^ CD8^−^) T cell phenotypes to DCLAMP^+^ cells by IF at varying distances.

